# A germline shortcut in protein language model retrieval of adaptive immune receptors

**DOI:** 10.64898/2026.07.21.739963

**Authors:** Suyue Wang, Ayan Sengupta, Songling Li, Daron M Standley

## Abstract

Protein language models (PLMs) are increasingly used to represent adaptive immune receptors; however, their advantages over classical alignment-based methods remain unclear. We benchmarked the ESM2 family, ESM-C, antibody-, and TCR-specific PLMs against sequence-alignment methods across four B-cell and T-cell receptor databases using CDR3, clonotype, paratope, and full variable domain representations. At the CDR3 level, alignments consistently out-performed all PLMs, except SCEPTR, a contrastively pre-trained TCR model that additionally encodes the germline V gene. BLOSUM62 and Levenshtein distance metrics exceeded all ESM2 variants by 7–12 percentage points in top-1 retrieval accuracy, and domain-specific PLMs did not close this gap. At the full variable domain level, the alignments and PLMs converged. Alignment performance peaked at the paratope level, whereas PLMs benefited primarily from the addition of conserved framework information in the full-length sequences. The one exception was heavily-mutated HIV-1 antibodies, where PLMs outperformed alignments. These results, together with germline reversion analyses, indicate that full-length PLM performance takes advantage of a germline shortcut rather than improved extraction of antigen-specific information from CDRs. We further showed that conventional random train-test splits inflate retrieval accuracy by 15–28 percentage points owing to clonal leakage. Together, these findings define the strengths and limitations of frozen zero-shot PLM embeddings for immune receptor retrieval and establish clone-aware benchmarking as a practical standard of comparison.

**Key Points:**

- Classical sequence alignment consistently outperformed frozen protein language models (PLMs) for CDR3 immune-receptor retrieval across the four datasets.
- Protein language models reached parity only in the full variable domain by exploiting the conserved germline-derived sequence, thus revealing a germline shortcut.
- Frozen PLMs exceeded alignment only for highly mutated HIV-1 antibodies.
- One PLM matching alignment at CDR3, SCEPTR, encoded germline V gene information.
- Random train–test splitting inflated accuracy by 15–28 percentage points; clone-aware splitting should be the standard.

**Bibliographical Note:** Daron M. Standley is Professor of Genome Informatics at the Research Institute for Microbial Diseases, Osaka University. His research developed computational methods for immune repertoire analysis and antibody discovery.

## 1 Introduction

Adaptive immunity relies on two classes of receptors to recognize antigens: B-cell receptors (BCRs, antibodies) and T-cell receptors (TCRs). Identifying receptors that recognize the same antigen or epitope is a central challenge in computational immunology and has applications in vaccine design, therapeutic antibody discovery, immune monitoring, and repertoire-based disease diagnostics.

A fundamental task is *antigen-specific receptor retrieval*; given a query receptor sequence, reference receptors known to bind the same antigen are identified. We evaluated retrieval using recall @1 (R@1), the fraction of queries whose top-ranked match shares the query’s antigen or epitope label. Performance depends on both the similarity metric and sequence representation, which range from the third complementarity-determining region (CDR3) to the full variable domain.

Traditional methods compare receptor sequences by alignment using BLOSUM62 (Henikoff and Henikoff, 1992) or edit distances. In contrast, protein language models (PLMs) encode sequences as embeddings compared to vector similarity. However, immune receptors differ from typical proteins. Their highly variable CDR3 is generated by V(D)J recombination with junctional insertions and deletions, and BCRs are further diversified by somatic hypermutation (SHM). Whether PLMs trained largely on evolutionarily related proteins can faithfully represent this non-evolutionary diversity remains unclear.

Recent studies raised similar concerns. Germline bias has been reported across antibody-specific and, to a lesser extent, general PLMs (Olsen et al., 2024). Zero-shot PLM embeddings also underperform handcrafted representations or sequence alignment for TCR specificity, unless retrained with TCR-specific objectives (Feng et al., 2025; Nagano et al., 2025; Frolenkova et al., 2025). In parallel, data leakage has been shown to affect the performance of protein-interaction and immune-receptor benchmarks (Bushuiev et al., 2024; Castorina et al., 2025).

Therefore, we asked the following three questions. Do frozen zero-shot PLMs outperform sequence alignment for antigen-specific retrieval? How does this comparison depend on the receptor representation? How should train–test splitting account for receptor similarity to avoid inflated performance?

To answer these questions, we benchmarked widely used PLMs, including ESM2, ESM-C, and receptor-specific models, against classical alignment across four immune-receptor databases spanning both BCRs and TCRs and a hierarchy of representations from the CDR3 to the full variable domain.

Our results revealed a consistent pattern across receptor types and databases. Alignment outperforms every general zero-shot PLM at CDR3, whereas the two approaches converge in the full variable domain. We showed that this convergence reflects a *germline shortcut* rather than an improved representation of antigen-binding features. Zero-shot PLMs provide additional signals, mainly for heavily hypermutated HIV-1 envelope antibodies. We further showed that clone-unaware train–test splitting inflates retrieval accuracy by 15–28 percentage points (pp), motivating clone-aware evaluation as a minimum reporting standard.

Our study established a systematic zero-shot benchmark across both receptor classes and multiple receptor representations. Unlike previous work focusing on supervised CDR3 embeddings or contrastive training, we characterize the frozen baseline against which such methods are compared and show that its apparent success at full length largely reflects germline information rather than the improved modeling of antigen-binding regions.

## 2 Materials and methods

We benchmarked sequence-alignment methods against PLM embeddings for antigen-specific receptor retrieval across four immune receptor databases and five receptor representations using clone-aware data splitting.

### 2.1 Benchmark overview

The evaluation followed a nearest-neighbor retrieval paradigm. For each seed (§2.5), the receptor sequences were split into a reference database (∼80%) and query set (∼20%). Each method ranked all reference sequences for every query by similarity, and retrieval was evaluated using recall @1 (R@1).

Top-score ties, common for CDR3 retrieval because identical CDR3s produce identical alignment scores and embeddings, were resolved by recall @1 (fractional credit equal to the proportion of tied candidates sharing the query label), making the metric deterministic. The tie statistics and sensitivity analyses are reported in the Supplementary Methods section.

### 2.2 Datasets and antigen labels

We used four publicly available immune receptor databases, annotated with antigen or epitope labels. All datasets underwent the same processing pipeline: removal of short or non-standard sequences, antigen name standardization, retention of the most frequent labels, and clone-aware 80:20 splitting repeated over 20 random seeds (§2.5). Distinct receptors sharing identical CDR3 were retained as separate entries, with the resulting ties handled as described above. The dataset compositions are summarized in Supplementary Table S2, and the processing details are provided in the Supplementary Methods.

IEDB (Vita et al., 2019) contributed antibody heavy chains under the ten most frequent protein antigens (∼680 sequences/seed). Sequences were renumbered with ANARCI (Dunbar and Deane, 2016) and trimmed to the variable domain. VDJdb (Goncharov et al., 2022) contributed to TCRβ CDR3s under ten epitope labels (∼1,000/seed). SAbDab (Dunbar et al., 2014) contributed paired antibodies under the five most frequent antigen labels (approximately 310/seed). McPAS-TCR (Tickotsky et al., 2017) contributed TCRs under ten epitope labels (∼1,000/seed).

Full-length TCRβ sequences were reconstructed by grafting CDR3s onto IMGT germline V/J templates (Lefranc et al., 2009); corresponding full-variable-domain analyses were therefore interpreted as directional. Download dates, label lists, and training and test set sizes are provided in the Supplementary Methods.

To assess possible pretraining leakage, BCR CDR3s were searched against UniRef90 with BLAST; seven sequences matched known broadly neutralizing antibodies but leave-one-out analysis showed a negligible effect on ESM2 performance (§3.4, Supplementary Note S2.3).

### 2.3 Receptor representations

We evaluated five receptor representations: CDR3, paratope, full variable domain, VJ clonotype, and V clonotype.

The first three progressively increase in sequence context, from CDR3 alone (the most variable receptor region and principal determinant of antigen specificity (Xu and Davis, 2000; Glanville et al., 2017)) to the paratope (CDR1–3, the antigen-binding loops (Liberis et al., 2018)) and the full variable domain (∼120 aa, including framework regions FW1–FW4). The latter two retain CDR3 but restrict retrieval to receptors sharing the same V/J genes (VJ-clonotype) or the V gene alone (V-clonotype), the standard clonotype definition (Bolotin et al., 2015); queries lacking a matching reference receive an R@1 of zero.

V/J assignments were obtained from database annotations (IEDB/SAbDab: ANARCI-assigned; VDJdb/McPAS-TCR: deposited annotations) and CDR3 boundaries followed IMGT numbering (Lefranc et al., 2009; Dunbar and Deane, 2016). Supplementary Table S12 summarizes the information obtained from each representation.

### 2.4 Methods compared

We compared sequence-alignment methods (Levenshtein distance, Smith–Waterman alignment with BLOSUM62, and BLAST), domain-specific metric TCRdist, and PLMs.

Alignment scores were computed with `parasail` (Daily, 2016), normalized by the longer sequence; gap parameters are given in the Supplementary Methods. BLAST (Altschul et al., 1990) confirmed that the alignment advantage was not specific to BLOSUM62 (Supplementary Table S10).

TCRdist (Dash et al., 2017; Mayer-Blackwell et al., 2021) was computed with `tcrdist3` using IMGT-defined CDR1, CDR2, CDR2.5 (the germline, V-gene-encoded loop between CDR2 and CDR3) and CDR3 distances from paired alpha–beta TCRs.

The PLMs included ESM2-150M, ESM2-650M, ESM2-3B (Lin et al., 2023), ESM-C-300M (EvolutionaryScale Team, 2024), AntiBERTy (Ruffolo et al., 2021), AbLang (Olsen et al., 2022) and TCR-BERT (Wu et al., 2021). Unless otherwise stated, embeddings were extracted from the final hidden layer without task-specific training (zero-shot), mean-pooled, and compared using cosine similarity. The CLS pooling results are reported in the Supplementary Material.

SCEPTR (Nagano et al., 2025), which jointly encodes CDR3 and V-gene information through contrastive pre-training, was evaluated separately (§3.5).

### 2.5 Clone-aware splitting, seeds and statistical analysis

To prevent clonal leakage, a recognized failure mode in which leakage-aware data splitting is designed to prevent (Joeres et al., 2025), receptors whose CDR3s shared ≥ 95% normalized Levenshtein similarity were clustered into single-linkage connected components, which were assigned entirely to the reference (∼80%) or query (∼20%) partition. Splitting was repeated over 20 random seeds, and the reported R@1 values were the seed means. Because the underlying sequence set remained fixed, the variation across seeds reflects train–test partitioning rather than biological sampling. The 95% threshold is a practical proxy for clonal relatedness; therefore, robustness was assessed across thresholds from 0.85 0.99 (§3.5).

Confidence intervals were estimated using nested bootstrap over seeds and queries (1,000 replicates) (Tsamardinos et al., 2018). Effect sizes were quantified using Hedges’ *g*, and practical equivalence for BCR full-variable-domain comparisons was tested using two one-sided tests (TOST) (Lakens, 2017) with Δ = 0.05 R@1. Recall@5 and the mean reciprocal rank (MRR) produced the same ranking. Additional statistical details are provided in Supplementary Methods and Supplementary Table S13.

## 3 Results

We first show that clone-aware splitting is necessary for unbiased evaluation (Section 3.1) and then compare alignment and PLMs across a representation ladder from CDR3 alone to the full variable domain. Alignment leads to CDR3 (§3.2), the two method families converge at full length through a germline shortcut (§3.3), and PLMs lead only to receptors that diverge strongly from the germline (§3.4). Section 3.5 extends the analysis to TCR-specific methods, paired-chain retrieval, and robustness.

### 3.1 Clone-aware splitting is a prerequisite

First, we quantified the bias introduced by conventional random train–test splitting. Relative to clone-aware splitting, random stratified splits substantially inflate retrieval performance (Figure 1). In a matched 20-seed comparison, random splitting increased ESM2-150M BCR Recall@1 from 0.353 to 0.632 at CDR3 (+27.9 pp) and from 0.507 to 0.674 in the full variable domain (+16.6 pp). Similar inflation occurred for the alignment methods (BLOSUM62: +20.4 and +18.7 pp, respectively). Across methods and representations, inflation ranged from ∼15 to 28 pp, indicating that it reflects clonal leakage rather than method-specific behavior. Therefore, we use clone-aware splitting and recommend it as a minimum standard for immune receptor retrieval benchmarks.

**Figure 1.**
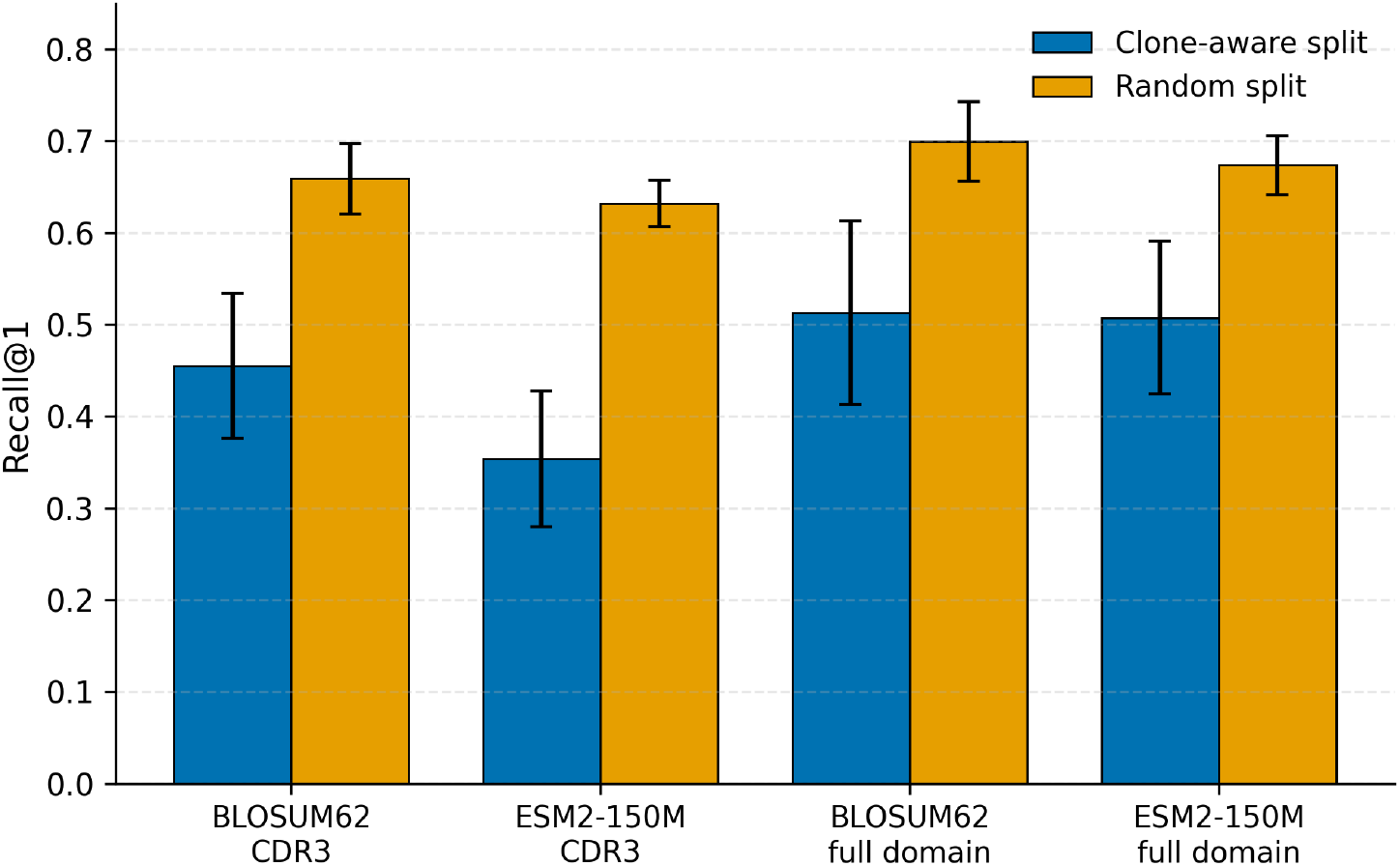
Random splits inflate immune receptor retrieval performance. Matched 20- seed BCR comparison: random stratified splits overestimate Recall@1 relative to clone-aware splits at both CDR3-only (CDR3) and full variable domain representations, for both alignment (BLOSUM62) and the PLM (ESM2-150M), a method-general artifact, not specific to PLMs (per-method inflation in §3.1). Error bars: SD across 20 seeds.

### 3.2 Alignment outperforms PLMs on CDR3-only retrieval

On CDR3-only inputs, the classical sequence comparison achieved the highest retrieval across all four datasets (Table 1; Figure 2). BLOSUM62 exceeded ESM2-150M (the smallest and most widely used ESM2 model, used here as the primary PLM comparator) by 10.2 pp on BCR, 9.4 pp on VDJdb TCR, 7.2 pp on SAbDab and 11.5 pp on McPAS-TCR. Levenshtein closely tracked BLOSUM62, and no ESM2 variant (150M–3B) surpassed either alignment method; even the best ESM2 model remained 6.8–8.7 pp behind. An independent BLAST+ baseline (`blastp-short`) yielded the same conclusion, ranking just below the best alignment method (0.6–2.9 pp) while outperforming every ESM2 variant (Supplementary Table S10). These gaps exceeded each dataset’s minimum detectable effect (∼2–5.5 pp recall @1 at 80% power), ruling out inadequate statistical power as an explanation.

**Figure 2.**
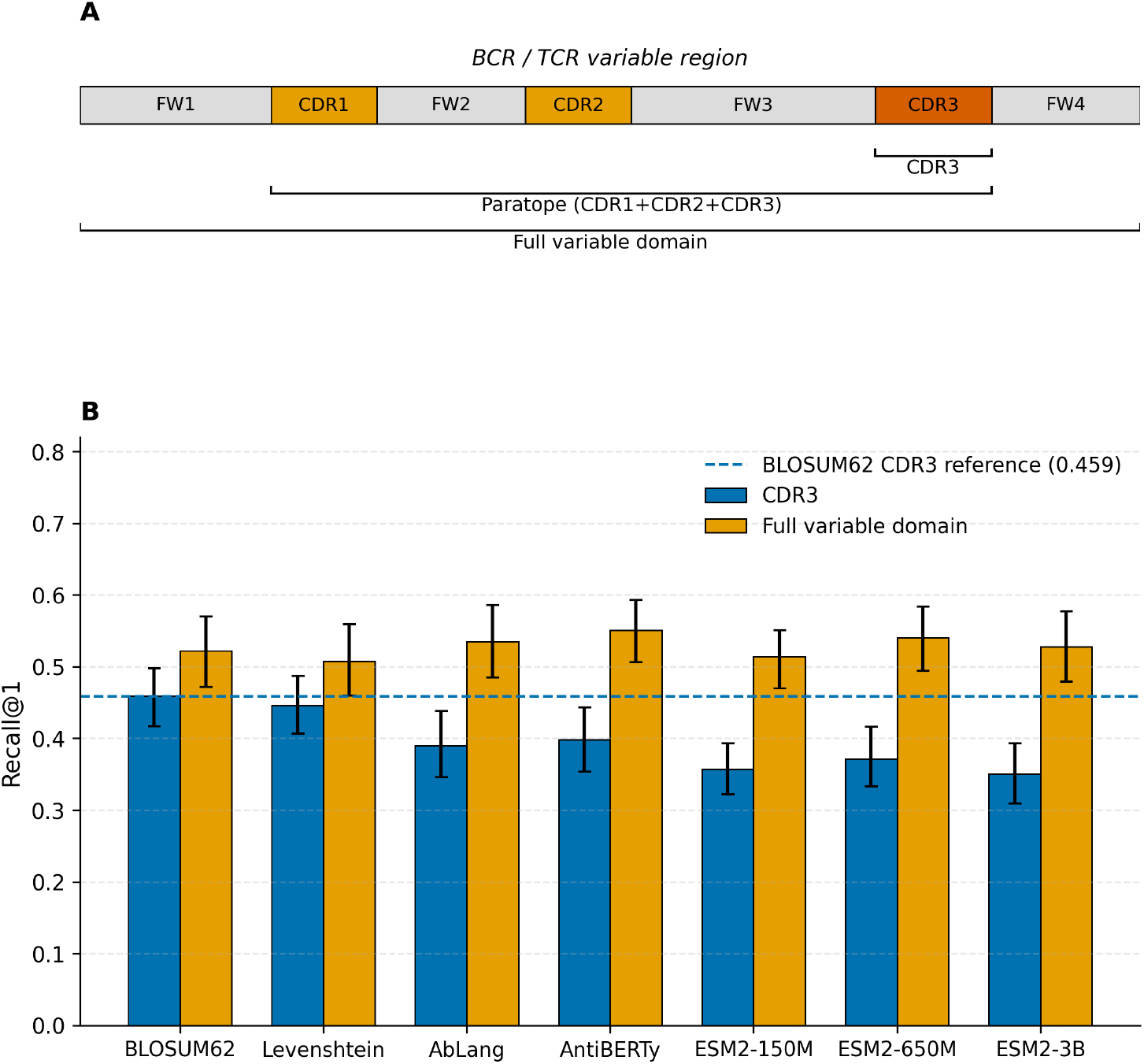
Benchmark design and BCR retrieval by representation. ***(A)*** Schematic of the receptor variable region (BCR heavy chain or TCR beta chain) with CDR3, paratope (CDR1 + CDR2 + CDR3) and full variable domain representation brackets. The clone-aware split groups sequences into connected components at CDR3 similarity ≥ 0.95. **(B)** BCR retrieval for alignment methods (BLOSUM62, Levenshtein), antibody-specific PLMs (AbLang, AntiBERTy) and the ESM2 family at the CDR3-only (blue) and full variable domain (orange) representations (the all-CDR paratope input, where alignment peaks, is in Table 1 and Figure 3); dashed line, 20-seed BLOSUM62 CDR3 reference (R@1 = 0.459). At the CDR3 the two alignment methods lead all PLMs; at the full variable domain several methods converge near R@1 ≈ 0.51–0.55 (including the antibody-specific AbLang 0.535 and AntiBERTy 0.551). Error bars: 95% nested-bootstrap CI across 20 seeds.

**Table 1.** Main zero-shot retrieval Recall@1. Two alignment methods (Levenshtein, BLOSUM62) and three ESM2 variants (150M, 650M, 3B). Bold marks the highest value in each row. Values are 20-seed nested-bootstrap means of expected R@1 (order-independent tie handling; see §2.1). BCR is shown at the CDR3 input, the all-CDR paratope, and the full variable domain; the single highest BCR value in the table is an alignment method at the paratope (Levenshtein 0.550, the antigen-binding loops with no framework). At the BCR full variable domain the five methods are close (R@1 range 0.508– 0.541, ∼3.3 pp spread), most pairs TOST-equivalent at Δ = 0.05 with the best-performing ESM2 (ESM2-650M) edging marginally above the weakest alignment method; see §3.3). VDJdb TCR, SAbDab and McPAS-TCR are shown at CDR3 as cross-database CDR3-level replication; their full representation ladders (CDR3 to full variable domain) are in Supplementary Table S1, and the paired-chain TCR ladder, including the purpose-built TCRdist (R@1 = 0.515, significantly above every other method), is in Table 2 (§3.5). Domain-specific PLMs (AntiBERTy, AbLang, TCR-BERT) and ESM-C-300M are reported in Supplementary Table S1.

| Dataset | Representation | Levenshtein | BLOSUM62 | ESM2-150M | ESM2-650M | ESM2-3B |
| --- | --- | --- | --- | --- | --- | --- |
| BCR (IEDB) | CDR3 | 0.446 | **0.459** | 0.357 | 0.372 | 0.350 |
|  | Paratope (all CDRs) | **0.550** | 0.538 | 0.445 | 0.417 | 0.446 |
|  | Full variable domain | 0.508 | 0.522 | 0.514 | **0.541** | 0.528 |
| VDJdb TCR | CDR3 | 0.355 | **0.375** | 0.282 | 0.304 | 0.307 |
| SAbDab | CDR3 | 0.351 | **0.358** | 0.286 | 0.273 | 0.259 |
| McPAS-TCR | CDR3 | 0.360 | **0.387** | 0.272 | 0.282 | 0.305 |

Effect sizes were uniformly large (paired Hedges’ *g* 1.04–3.72 across datasets), and the ranking was unchanged under every tie-handling convention (Supplementary Methods).

Model scale, domain specialization, and newer architecture did not close the gap. No ESM2 model matched alignment on any dataset; the antibody-specific AntiBERTy (0.398 at BCR CDR3, 6.1 pp below BLOSUM62) and TCR-BERT likewise fell short, and the newer ESM-C-300M performed similarly to ESM2-150M, remaining 7.1–9.9 pp below BLOSUM62 across datasets (Supplementary Table S15).

The deficit is also a property of embeddings, rather than a retrieval metric. A linear probe trained on frozen CDR3 embeddings likewise failed to match alignment, correcting embedding anisotropy (the embeddings’ skewed geometry) improved Recall@1 by at most 3–5 pp, which persisted under an alternative distance metric and disappeared for full-variable-domain embeddings despite their similar geometry (Supplementary Note S2.6; Figure S11). The deficit was largest for the CDR3s that ESM2 considered most unusual, yet did not track divergence from the germline, leaving the antigen-discriminating feature missed by the embeddings unresolved.

One clear exception, the most heavily hypermutated antibody, is examined in Section 3.4.

### 3.3 At full length, PLMs and alignment converge through a germline shortcut

In the full-variable-domain representation, the CDR3 performance hierarchy disappears. The five primary benchmark methods (two alignment methods and three ESM2 models) fell within a ∼3.3-pp range (0.508–0.541; Table 1), with overlapping 95% confidence intervals. Practical equivalence testing (TOST; §2.5) placed most comparisons within the prespecified Δ = 0.05 Recall@1 margin, including the primary comparison between ESM2-150M and BLOSUM62 (mean difference -0.005 across seeds). The only exception was ESM2-650M versus levenshtein (+0.034), which only marginally exceeded the equivalence margin. Therefore, the CDR3 ordering was effectively reversed, with the best-performing ESM2 model edging slightly above the weaker alignment baseline (Figure 2; Supplementary Table S4).

The representation ladder shows how this convergence occurred (Figure 3). At the CDR3, alignment exceeded ESM2-150M by 10.2 pp. The clonotype representations (VJ-clonotype and V-clonotype) mainly altered whether a query had an eligible reference rather than how candidates were ranked: the stricter VJ filter drove every method to a common floor by eliminating many valid matches, whereas the V-only filter improved retrieval where coverage was permitted (IEDB BCR and McPAS-TCR, but not VDJdb). The alignment maintained a modest advantage throughout these intermediate representations (Supplementary Table S1).

**Figure 3.**
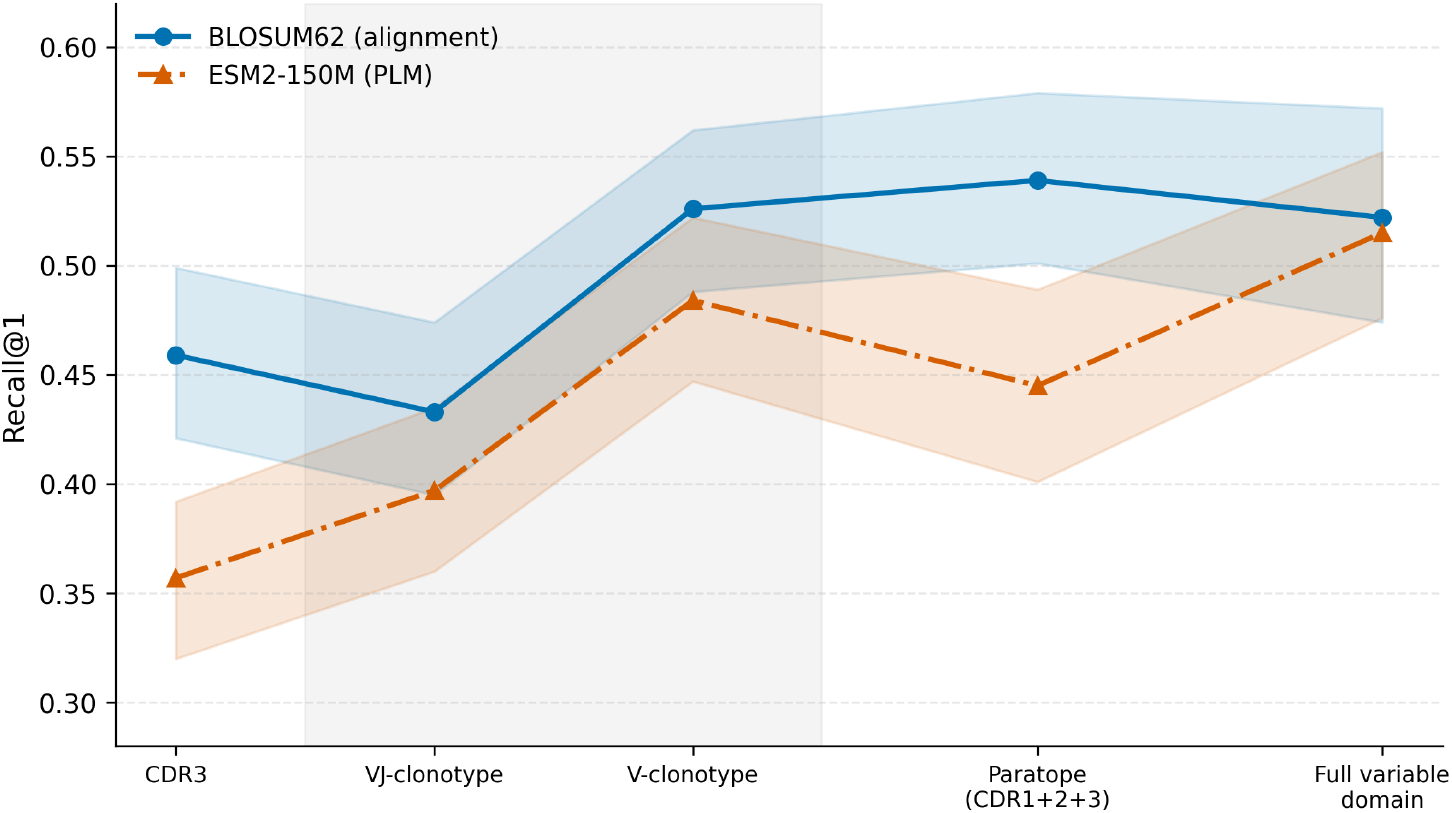
BCR retrieval across the representation ladder. BCR Recall@1 across the ladder in defined order (CDR3, VJ-clonotype, V-clonotype, paratope and full variable domain; 20-seed nested bootstrap). The gray band marks the two clonotype-retrieval variants (VJ-clonotype, V+J; V-clonotype, V only), which keep the CDR3 input but add a germline-gene filter (§2.3); the other representations vary input content. These are two distinct axes (input content vs. retrieval constraint) and should not be read as a single monotonic trend along the x-axis. BLOSUM62 (blue) leads at the CDR3 input and peaks at the paratope, gaining nothing from the framework; ESM2-150M (orange) trails alignment from the CDR3 to the paratope and reaches parity only at the full variable domain (germline-reversion analysis in §3.3; per-representation values with 95% CIs in Supplementary Table S3). Shaded bands: 95% nested-bootstrap CI.

Adding CDR1 and CDR2 to form the complete paratope improved retrieval for both method families by approximately 8 pp, but PLMs still trailed alignment by 9–13 pp. Only after the conserved framework was added did their trajectories diverge; PLMs improved further, whereas alignment declined slightly, bringing the two approaches to practical equivalence. Therefore, alignment peaks at the paratope, whereas PLMs reach parity only in the full variable domain.

This pattern localizes the antigen-discriminating signal to the CDRs. Alignment achieved its highest performance on the paratope (the best BCR result in Table 1, Levenshtein R@1 = 0.550), and adding the conserved framework slightly reduced performance because the framework diluted CDR differences without contributing additional antigen-discriminating information. (Part of the gain from CDR3 to the paratope is germline-derived through CDR1 and CDR2, as confirmed below.) In contrast, every PLM, including AntiBERTy and AbLang (Supplementary Table S15), benefited from the framework, which is consistent with the reported germline bias of antibody language models (Olsen et al., 2024).

Only PLMs have benefited from this framework. The method × representation interaction was positive in all 24 method × model size × dataset comparisons (paired Hedges’ *g* 0.60– 1.95). Because these comparisons are not independent, we treated the four databases rather than the individual cells as the unit of replication; all four showed the same interaction (Supplementary Note S2.2). However, this interaction does not establish a germline shortcut because the framework could, in principle, contain antigen-discriminating information accessible only to PLMs. The following analyses distinguish between these possibilities:

Therefore, we refer to this convergence as a *germline shortcut*: zero-shot PLMs close their CDR3 deficit by exploiting the conserved germline sequence in the framework, whereas alignment, which is already saturated on the CDRs, does not benefit from it and declines slightly. This does not imply that full-variable-domain retrieval is determined solely by germline identity because full-variable-domain performance substantially exceeds that of a V-gene-only baseline (below). The same conclusion was obtained using alternative embedding-pooling strategies (Supplementary Note S2.2).

#### Germline-reversion test

As a direct test, we replaced the antibody framework with its inferred germline sequence while retaining the native CDRs and separately replaced CDR1 and CDR2 with their germline sequences while retaining the native framework (20 paired seeds). For ESM2-150M, the germline framework alone accounted for approximately two-thirds of the PLM’s improvement from the paratope to the full variable domain (P < 10⁻⁵). Replacing CDR1 and CDR2 with germline sequences produced no detectable loss, and framework somatic mutations contributed little overall (with no measurable benefit outside HIV-1 antibodies; Section 3.4). In contrast, BLOSUM62 levels remained essentially unchanged throughout the experiment. The same pattern was reproduced for the independent SAbDab dataset (Supplementary Note S2.5). This experiment provided the strongest evidence that full-variable-domain convergence arises through a germline shortcut.

#### V-gene oracle

Next, we asked whether the observed convergence could be explained simply by V-gene identity. Across all full-variable-domain methods, 91.5–99.2% of top-ranked retrievals shared the query’s V-gene family, far above the random expectation (∼34%; Supplementary Figure S1, Table S11, and Note S2.4). However, explicitly providing the correct V-gene family did not improve retrieval. A two-stage oracle (true V-gene family followed by full-variable-domain BLOSUM62) achieved R@1 = 0.507 (95% CI, 0.464– 0.550), essentially identical to full-variable-domain BLOSUM62 alone (0.513 on the V-assignable subset), whereas a V-gene-only baseline reached only 0.190 (95% CI, 0.181– 0.200). Full-variable-domain alignment, therefore, captures essentially all retrieval signals associated with V-gene identity, whereas V-gene information alone is far from sufficient for accurate retrieval. We regard the oracle as a consistency check rather than decisive evidence, with the germline reversion experiment carrying the primary weight.

Each analysis has limitations (Supplementary Note S2.2); therefore, we present the germline shortcut as a working model rather than a proven mechanism. Nevertheless, it is supported by three independent observations that do not rely on assignable V genes: germline-V enrichment in native TCR retrieval, loss of SCEPTR’s apparent advantage after clonotype filtering (§3.5), and embedding-geometry controls showing that the CDR3 deficit reflects the embeddings themselves rather than the similarity metric (Supplementary Note S2.6).

### 3.4 A directional exception: PLMs lead on the most hypermutated antibodies

The overall convergence of the full variable domain concealed a clear antigen-specific exception (Supplementary Figure S7; Table S8). BLOSUM62 outperformed ESM2-150M for most antigens, including *Plasmodium* CSP, SARS-CoV-2 spike, and influenza HA (by 7–10 pp), whereas the HIV-1 envelope showed the opposite pattern, with ESM2 leading by 21.2 pp. Therefore, HIV-1 was the only major antigen for which zero-shot PLM clearly outperformed sequence alignment.

Because broadly neutralizing HIV-1 antibodies are extensively mutated and well-represented in public databases, we tested whether this advantage could reflect pretraining leakage. A BLAST audit identified seven CDR3s matching documented broadly neutralizing antibodies in UniRef90, but excluding them changed the ESM2 advantage by only +1.1 pp. The leak-free HIV-1 subset still favored ESM2 by 22.2 pp (Supplementary Note S2.3), arguing against memorization as an explanation.

Stratifying antibodies by SHM was consistent with an association between germline divergence and PLM performance but did not support a general SHM-dependent mechanism. ESM2 led by 7.0 pp in the highest-SHM bin; however, this crossover was driven largely by the HIV-1 envelope, disappearing within influenza HA and other non-HIV antigens at comparable SHM levels (Supplementary Note S2.3; Table S9). Therefore, we interpret this as a directional, hypothesis-generating exception, rather than evidence for a general SHM mechanism. The present data support only an association between strong germline divergence and improved zero-shot PLM performance, and not the mechanism underlying this advantage (see also Supplementary Note S2.5).

### 3.5 Generality and robustness

#### TCR and paired chains

The TCR representation ladder mirrored the BCR patterns (Supplementary Table S1). Because full-length TCRβ chains were reconstructed from germline templates (§2.2), the full-variable-domain TCR results serve primarily as a constructive control; the mechanistic interpretation instead rests on the clonotype-filter convergence and native-TCR V-gene enrichment. Paired VDJdb analysis showed the same pattern (Table 2). Alignment peaked again at the paratope, whereas ESM2 reached parity only in the full variable domain. The paired clonotype filters sharpened this convergence further, requiring both alpha- and beta-chain V and J genes to match, leaving almost every query without an eligible reference, collapsing all methods to the same floor (paired VJ-clonotype R@1 = 0.122–0.123). Germline-associated retrieval was also evident in native TCRs without germline reconstructions. At the CDR3, correct top-ranked hits shared the query’s beta-chain variable (TRBV) gene family in 67–70% of cases for both alignment and ESM2 versus 7.8% expected by chance (∼9-fold enrichment). TCRdist remained the strongest performer (R@1 = 0.515 [0.489, 0.541]), benefiting most from paired chains (+8.7 pp versus +3.7 pp for BLOSUM62). Because it explicitly scores germline-derived CDR1, CDR2 and CDR2.5 together with CDR3, part of its advantage is germline-informed.

**Table 2.** TCR paired-chain ladder (20 seeds, VDJdb). Expected R@1 for five benchmark methods across the paired ladder in defined order (10 fixed epitope labels, clone-aware split). The input representations (CDR3, paratope and full variable domain) are interleaved with the two clonotype-retrieval variants (VJ-clonotype†, V+J on both chains; V-clonotype†, V on both chains), which keep the paired CDR3 input but add a germline-gene filter (§2.3). **Bold** marks each method’s highest value among the input representations. The full-variable-domain paired input uses germline-reconstructed full variable domains and is interpreted as directional (§2.2). tcrdist3 is a single value because tcrdist3 computes one IMGT-weighted distance over CDR1/CDR2/CDR2.5/CDR3 of both chains, independent of the input ladder; it is reported alongside the full-variable-domain paired result for direct comparison.

| Method | CDR3 paired | VJ-clonotype paired† | V-clonotype paired† | paratope paired | full variable domain paired |
| --- | --- | --- | --- | --- | --- |
| Levenshtein | 0.414 | 0.123 | 0.180 | **0.443** | 0.437 |
| BLOSUM62 | 0.436 | 0.123 | 0.179 | **0.459** | 0.437 |
| ESM2-150M | 0.352 | 0.123 | 0.178 | 0.395 | **0.424** |
| ESM2-650M | 0.349 | 0.122 | 0.178 | 0.396 | **0.427** |
| ESM2-3B | 0.363 | 0.123 | 0.180 | 0.399 | **0.430** |
| tcrdist3 | — | — | — | — | **0.515** |

† Clonotype variants (§2.3): Reference candidates are restricted to those sharing the query’s germline genes on *both* chains (V and J for VJ-clonotype; V only for V-clonotype), then ranked by the paired CDR3; a query with no same-clonotype reference is a miss. Requiring a paired match is strict, so VJ- and V-clonotype pairs fall far below CDR3; under either filter, all five methods are statistically indistinguishable (full discussion in §3.5; 95% CIs in paired\_clonotype\_20seed\_nested\_ci.csv).

#### Pairing

Antibody heavy–light pairing left all five methods statistically indistinguishable in both IEDB (R@1 0.52–0.54) and SAbDab pairs (0.35–0.36; Supplementary Table S5). Therefore, pairing behaved asymmetrically, substantially improving TCR retrieval (especially TCRdist) but contributing little to antibodies, an observation we note but do not explain, given the limited paired-antibody datasets (§4.2).

#### SCEPTR, a contrastively pre-trained TCR language model

The use of only PLM to match alignment also supports the germline shortcut rather than contradicting it. SCEPTR (Nagano et al., 2025) jointly encodes the V gene and CDR3 through a TCR-specific contrastive objective, rather than serving as a general zero-shot protein embedding. Under the same clone-aware benchmark and common gene-mappable query set, alignment-level CDR3 retrieval (McPAS-TCR R@1 = 0.457; VDJdb 0.333) matched TCRdist and exceeded BLOSUM62 by 2–6 pp, whereas the general ESM2 family remained 11–18 pp lower (Supplementary Table S14). This advantage is germline-associated, rather than antigen-specific. Under the same clonotype constraints applied to every method, SCEPTR lost its CDR3 advantage once retrieval was restricted to receptors sharing the query V (or V and J) gene, converging with or falling below BLOSUM62 (Supplementary Table S1). Even incorrect top-ranked matches shared the query’s V-gene family more often than alignment did, indicating that the representation strongly prioritizes germline identity. Similarly, on paired alpha–beta chains, SCEPTR converged with generic alignment (0.468 versus 0.467), leaving only TCRdist (0.490) ahead on the common evaluation subset. Thus, even the strongest task-specific PLM reaches the same germline-informed frontier, rather than recovering additional antigen-discriminating signals beyond alignment. Because SCEPTR’s original benchmark included VDJdb-derived data, McPAS-TCR provides more independent validation.

#### Robustness

The CDR3 advantage of alignment was robust to the clone definition and dataset composition. BLOSUM62 consistently outperformed ESM2-150M across CDR3 length strata (Supplementary Figure S3), the most evaluable BCR antigen classes (Supplementary Table S7), and epitope-level VDJdb labels. Varying the clone threshold from 0.85 to 0.99 likewise preserved both the CDR3 advantage (10–13 pp) and the practical equivalence at the full variable domain (Supplementary Figure S4; Table S6). Across receptor classes, databases, chain configurations, and clone definitions, the same qualitative pattern emerged: alignment extracted more antigen-relevant information from CDR3, whereas PLMs reached parity only after conserved germline information was introduced through the full variable domain. Therefore, the central conclusions were not specific to a particular receptor type, benchmark dataset, or analytical choice.

## 4 Discussion

Using a clone-aware benchmark spanning four immune receptor databases, we found that classical sequence alignment consistently outperformed frozen zero-shot PLMs for CDR3 retrieval, whereas the difference largely disappeared in the full variable domain. Together with the germline reversion and V-gene analyses, these findings support a germline shortcut rather than improved antigen-specific representations as the primary explanation for full-length convergence (Figure 4).

**Figure 4.**
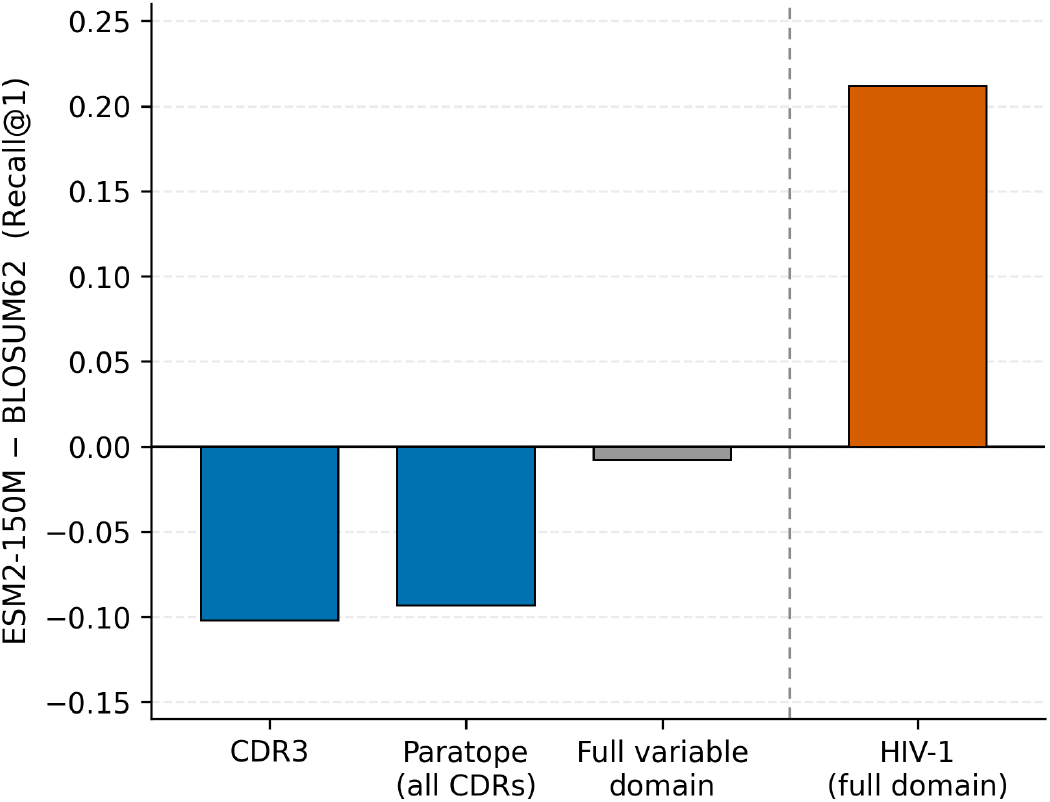
The germline axis in one view (BCR; illustrative). ESM2-150M minus BLOSUM62 Recall@1 across four settings grouped into three germline regimes; bars are colored by which family leads (alignment blue, parity gray, PLM orange). Same antibodies, increasing germline content of the input (left of the divider): with CDR-level input and no framework, alignment leads at CDR3 only (CDR3, -0.10) and at all CDRs (paratope, -0.09); adding the conserved germline framework at the full variable domain (-0.01, TOST-equivalent) closes the gap, the visual signature of the germline shortcut. Different antibodies, same full-variable-domain input (right of the divider): on the heavily hypermutated, far-from-germline HIV-1 envelope subset the PLM leads (full variable domain, +0.21; leak-free +0.22, §3.4). The first three bars vary receptor representation for one antibody set; the fourth changes the antibody set (not the representation), so it is offset. The regional axis (framework vs. CDR3) locates where the PLM trails, but the within-CDR3 mechanism is not captured by a model-independent germline distance and remains open (Supplementary Note S2.6). This panel summarizes the BCR ESM2-150M-vs-BLOSUM62 contrast; full cross-database and cross-size evidence is in Table 1 and §3.5. Values are 20-seed nested-bootstrap means (Table 1; §3.4, Supplementary Note S2.3).

We used germline shortcuts to describe a representational rather than a biological phenomenon. Germline genes are unquestionably important for antigen recognition, and specific V-gene families define well-characterized antibody classes (for example, VH1-69 in influenza (Lingwood et al., 2012) and VH1-2 in HIV-1 CD4-binding-site antibodies (Zhou et al., 2010)), but in our benchmark PLMs reached parity with alignment only after conserved germline-derived framework sequence became available. Germline-reversion and V-gene analyses point to the same interpretation, while even TCRdist and contrastively pre-trained SCEPTR derive part of their advantage from explicitly incorporating germline-derived information. The CDR3 deficit and full-length convergence therefore appear to reflect the same representational bias: frozen embeddings capture conserved germline contexts more effectively than the diverse junctional features that distinguish antigen specificities.

The principal exception was HIV-1 envelope antibodies, the only major antigen class for which PLMs substantially outperformed alignment. These antibodies are among the most highly mutated regions of the antibody sequence space (Burton and Hangartner, 2016). Although our data do not establish a causal role for somatic hypermutation, the exception occurs where receptors diverge furthest from the germline, suggesting that zero-shot PLMs may be most useful for highly evolved antibody lineages rather than for typical repertoires. As somatic hypermutation is unique to B cells, this regime is not expected for TCRs.

Our conclusions apply specifically to frozen zero-shot embedding. This does not imply that PLMs are intrinsically unsuitable for immune receptor analysis. Fine-tuning, contrastive learning, or explicit incorporation of germline information may outperform alignment; recent studies support this possibility (Nagano et al., 2025; Wang et al., 2024, 2025). Instead, our contribution is to establish a zero-shot baseline against which such methods should be evaluated and to explain why full-length convergence can occur despite inferior CDR3 retrieval.

More broadly, these findings illustrate a recurring challenge in biological machine learning: a strong benchmark performance does not necessarily imply that a model has learned the intended biological signal. In immune-receptor retrieval, near-parity at full length can arise largely from germline-associated information rather than from improved encoding of antigen-discriminating features. Distinguishing between these signal sources will therefore be important both for evaluating future immune-receptor representations and for understanding what protein language models encode in a zero-shot setting.

### 4.1 Practical implications

These findings motivated six recommendations for immune-receptor retrieval benchmarks (Supplementary Box S1): use clone-aware splitting, report tie-aware evaluation metrics, include a zero-shot alignment baseline, stratify performance by receptor representation and germline context, assess robustness across clone definitions, and report embedding provenance.

In practice, the recommendations are straightforward (Table 3). For CDR3-only retrieval, BLOSUM62 or Levenshtein remains the default zero-shot method. For BCRs, alignment on the complete paratope provided the highest retrieval accuracy, whereas adding the framework did not further improve the alignment. In the full variable domain, ESM2-150M and BLOSUM62 performed similarly overall, so antigen composition may guide the method choice. The TCRdist remains the strongest method for paired TCR retrieval. Random train– test splitting should be avoided because it substantially affects the retrieval performance.

**Table 3.** Practical method recommendations.

| Setting | Recommended zero-shot method | Rationale |
| --- | --- | --- |
| CDR3-only<br>BCR/TCR retrieval | BLOSUM62 or<br>Levenshtein | Consistent CDR3 advantage over PLMs |
| Best zero-shot accuracy (BCR) | BLOSUM62 or Levenshtein on all CDRs (paratope) | Highest R@1 overall; the framework (full variable domain) does not help alignment |
| Full variable domain BCR, mean-pooled ESM2 available | BLOSUM62 or ESM2-150M | Comparable aggregate R@1; inspect antigen and germline strata |
| Paired TCR chains | TCRdist (tcrdist3) | TCRdist (R@1 = 0.515) significantly exceeds every tested alignment and ESM2 variant (R@1 0.42–0.44); BLOSUM62, Levenshtein and the three ESM2 sizes are mutually comparable |
| Any benchmark evaluation | Clone-aware splitting | Random splits inflate R@1 by 15–28 pp |

Recommendation Rank by Accuracy. The retrieval time cost differs and partly offsets the gap: pairwise alignment recomputes a score for every query–reference pair, whereas PLM embeddings are computed once and retrieval is a vector-similarity lookup. Thus, on very large reference banks, the embedding route can be markedly faster despite its lower accuracy.

Although alignment generally provides the highest zero-shot accuracy, PLM-based retrieval has an important computational advantage: embeddings are computed once and reused, whereas alignment requires pairwise comparisons for every query–reference pair. Embedding-based retrieval may therefore remain preferable for very large reference databases, where the search speed outweighs modest differences in retrieval accuracy.

### 4.2 Limitations

This study has several limitations.

First, the available antigen label space is limited. Each benchmark contained only 5–10 antigen or epitope labels, and protein-level BCR annotations may group antibodies targeting distinct epitopes. Although the principal findings were replicated across both protein- and peptide-level datasets, broader epitope-resolved benchmarks would better distinguish germline-associated from antigen-specific retrieval.

Second, the apparent PLM advantage among highly mutated antibodies is largely driven by the HIV-1 envelope. As additional highly mutated antigens are scarce, we could not separate the effects of somatic hypermutation from antigen identity.

Third, full-length TCRβ sequences were reconstructed from germline templates rather than being directly observed. Accordingly, the strongest mechanistic evidence for the germline shortcut comes from native antibody analyses, whereas TCR interpretation relies primarily on clonotype filtering, representation effects, and germline enrichment in native TCR retrieval.

Finally, our conclusions are limited to frozen, last-layer, and mean-pooled zero-shot embedding. We did not systematically evaluate alternative pooling strategies, intermediate layers, fine-tuning, contrastive learning, joint receptor–antigen models, or the rapidly expanding family of newer PLMs. Likewise, although robustness analyses suggest that neither alternative similarity metrics nor clone definitions would alter the principal conclusions, these remain methodological choices rather than biological ground truth.

### 4.3 Future directions

Future work should extend these analyses to larger epitope-resolved datasets, particularly to native paired full-length receptors and highly mutated antibodies beyond HIV-1. Germline-aware, contrastively trained, and fine-tuned PLMs should then be evaluated across the same representation hierarchy to determine whether they can overcome the CDR3 limitation observed in the zero-shot setting.

Fundamentally, our results narrow down the remaining questions. Because the CDR3 deficit cannot be explained by the similarity metric, embedding geometry, or simple divergence from the germline, the critical issue is which junctional features frozen PLMs fail to encode, and whether those features can be recovered through improved pretraining or task-specific adaptation.

## Acknowledgements

We thank the developers and maintainers of the IEDB, VDJdb, McPAS-TCR, and SAbDab for providing the public immune receptor datasets used in this benchmark.

## Ethics statement

This study used only publicly available, de-identified immune receptor sequences from IEDB, VDJdb, McPAS-TCR, and SAbDab; no new human subject data were collected. These datasets were originally gathered with institutional review board approval (see primary publications).

## Author contributions

CRediT taxonomy. S.W.: Conceptualization, Methodology, Software, Writing of the original draft. A.S.: Methodology, Software, Writing – Review, and Editing. S.L.: Formal analysis, Validation, Data curation, and visualization. D.M.S.: Supervision, Project administration, writing – review, and editing. All authors have reviewed and approved the final manuscript.

## Data availability

The benchmark framework, evaluation and analysis codes, and download scripts are available at https://github.com/NidRUAwake/immune-embedding-benchmark. The source datasets were obtained from their repositories under their licenses and were not redistributed; license-permitted processed inputs and MD5 checksums enabled reproduction were provided. Precomputed embeddings can be regenerated using the scripts provided. The model weights were obtained from HuggingFace or the corresponding Python packages.

