## Supplementary Materials for "A germline shortcut in protein language model retrieval of adaptive immune receptors"

#### 1 Supplementary Methods

This section gives extended technical details for the benchmark beyond the main paper Methods (§2): dataset construction, PLM embedding pipeline, alignment/distance scoring, and statistical analysis.

##### 1.1 Dataset construction

**IEDB BCR.** B-cell assay records were downloaded from IEDB (Vita *et al.*, 2019; release as cited in main Methods), filtered to human heavy-chain entries with paired antigen labels. Distinct receptor records that share an identical CDR3 were retained (so identical CDR3s recur in the bank, handled by the expected-R@1 tie convention, §Tie handling). Sequence cleaning removed records with non-canonical residues, premature stop codons, or CDR3 length outside the empirical range [5, 35] amino acids. Antigen labels were harmonized using a curated alias mapping (`label_aliases.json`) merging synonymous strings such as "SARS-CoV-2 spike" / "S protein [SARS-CoV-2]" / "Spike glycoprotein [SARS-CoV-2]". After filtering to the top 10 antigen labels by record count (`–top-labels 10 –min-label-count 14`), the working dataset contained approximately 675 sequences per seed (n_train ≈ 546, n_test ≈ 128 under the 80:20 clone-aware split). The antigen labels used are protein-level categories (e.g., "HIV-1 envelope glycoprotein", "Influenza A hemagglutinin", "*Plasmodium falciparum* CSP") rather than epitope-level peptide labels; this coarse labeling is a known limitation (§4.2 of the main text). The CDR3 amino-acid composition of this benchmark differs from the UniProt background in the expected V(D)J-junction-enriched manner (Supplementary Figure S5).

**Standardizing the antibody sequences.** IEDB deposits are not all the same kind of molecule: some entries are a single antibody heavy-chain variable region, many are that variable region joined to its constant region (a longer Fab-heavy fragment), and a small number are a heavy and a light chain joined into one string. Left unprocessed, this format heterogeneity allows retrieval to match sequences by shape rather than biology: for example, two antibodies that both carry the antigen-independent constant region resemble each other regardless of which antigen they bind, and we found this format correlated with antigen identity. To remove this artifact we re-numbered every IEDB antibody with ANARCI (an automated antigen-receptor numbering tool) under the IMGT (ImMunoGeneTics) numbering scheme and retained only the heavy-chain variable domain (the antigen-contacting region), discarding signal peptides and constant regions so that every full-length (full variable domain) input is the same kind of molecule (consistent with how SAbDab sequences are defined). From the same numbering we read out CDR1, CDR2, CDR3 and the V/J germline gene calls; the CDRs build the multi-CDR (paratope) input and the V/J calls provide the germline-gene filter used by the VJ- and V-clonotype retrieval (§2.3), because IEDB's own CDR1/CDR2 columns were empty for 77–90% of records and its J-gene column for ≈ 89%. Records ANARCI identified as paired heavy+light antibodies (197) were routed to a separate paired analysis, since a single chain cannot represent the complete paratope (the antigen-binding surface formed by the heavy and light chains together); light-chain-only, non-antibody, or otherwise unparseable records (103 total) were excluded. For T-cell receptors, where only CDR3 and V/J gene names are deposited (no full-length sequence to re-number), records lacking a V-gene (CDR3-only deposits; ≈ 10% of VDJdb, ≈ 5% of McPAS) were excluded from the multi-CDR representation rather than padded, as their CDR1/CDR2 cannot be determined.

**Antibody PLM weights.** AbLang and AntiBERTy embeddings were originally computed from HuggingFace weights, whose repositories became unavailable during revision. Both were recomputed with their published Python packages (identical models, bundled weights); they encode amino-acid sequences only. While recomputing AntiBERTy we found that the original embeddings had been generated without the per-residue (whitespace-separated) tokenization the model requires (tokenization is how a language model splits its input into the discrete units, or tokens, that it reads; these protein models expect one token per amino acid), so each CDR3 had collapsed to a single unknown token and the embeddings carried almost no sequence information. With the correct tokenization, AntiBERTy's BCR CDR3 Recall@1 rises from 0.261 to 0.398; we report the corrected values throughout. Because the VJ- and V-clonotype variants now rank by the CDR3 embedding (the gene only enters as a filter, §2.3), AbLang and AntiBERTy are assessable at VJ- and V-clonotype as well as at CDR3, and both are reported across the full representation ladder: the CDR3 (CDR3), the clonotype variants (VJ- and V-clonotype), the all-CDR input (paratope), and the full chain (full variable domain).

**VDJdb TCR.** Human VDJdb beta-chain records (Goncharov *et al.*, 2022) were taken from the VDJdb 2025-12-29 release (downloaded 2026-04-27 from the official GitHub release artifact `vdjdb-2025-12-29.zip`; subsequent releases (2026-05-16, 2026-06-03) are acknowledged but not used, to keep a fixed evaluation snapshot). The benchmark filters epitope labels by intersecting the top-15 most-abundant labels with those having ≥ 50 records at every representation (CDR3 to the full variable domain); five epitopes are excluded by this intersection because they lack V/J annotation needed for representations beyond CDR3, leaving 10 working labels (`SLLMWITQV`, `NLVPMVATV`, `KLGGALQAK`, `GILGFVFTL`, `VISNDVCAQV`, `GLCTLVAML`, `RAKFKQLL`, `ELAGIGILTV`, `AVFDRKSDAK`, `YLQPRTFLL`). Per-label capping (`–max-per-label-tcr 100`) gives ≈ 1,000 sequences per seed, with ≈ 801 in train and ≈ 199 in test under the 80:20 clone-aware split. CDR3 sequences were extracted under IMGT conventions; conserved cysteine (104C) and phenylalanine/tryptophan (118F/W/Y) flanking residues were stripped from the CDR3 input. The VJ- and V-clonotype variants restrict candidates to the query's TRBV gene (and TRBJ for VJ-clonotype), matched at gene level, then rank by CDR3 (§2.3). The canonical TCR CDR3 evaluation used 20 seeds.

**SAbDab.** Human antibody structure records were downloaded from SAbDab (Dunbar *et al.*, 2014). Records with complete CDR annotation (CDR1, CDR2, CDR3 for both heavy and light chain) and a paired antigen identifier were retained. Filtering to the top 5 antigen labels with at least 25 records each (`–top-labels 5 –min-label-count 25 –max-per-label-sabdab 100`) yielded a working dataset of ≈ 310 sequences per seed (n_train ≈ 251, n_test ≈ 59). SAbDab uses the same 20 seeds as the other databases; its wider confidence intervals reflect the smaller working dataset (n_test ≈ 59). Because per-seed SAbDab test sets vary widely in size, the query-weighted CDR3 R@1 (0.362, pooled over all test queries through the nested bootstrap, as reported in main-text Table 1) differs from the unweighted per-seed mean (near 0.43); both aggregations yield the same alignment-over-ESM2 ordering.

**McPAS-TCR.** McPAS-TCR (Tickotsky *et al.*, 2017) records were filtered to the top 10 most-abundant epitope labels with at least 30 records each at every representation (GILGFVFTL, GLCTLVAML, NLVPMVATV, LPRRSGAAGA, WEDLFCDESLSSPEPPSSSE, CRVLCCYVL, EAAGIGILTV, RFYKTLRAEQASQ, TPRVTGGGAM and VTEHDTLLY), with `–max-per-label-tcr 100` for evaluation parity with the VDJdb TCR benchmark (which likewise resolves to 10 working labels). The canonical McPAS CDR3 evaluation used 20 seeds with n_train ≈ 801, n_test ≈ 199 per seed. CDR3 sequences underwent the same preprocessing as VDJdb (104C and 118F/W/Y anchors stripped). Full-length TCRβ sequences were reconstructed from IMGT germline templates by concatenating the framework regions (FW1–FW4) with the called V and J segments. Reconstructed sequences are interpreted as directional rather than definitive evidence for full-length performance. *Epitope overlap with VDJdb.* Three of the McPAS labels (GILGFVFTL, GLCTLVAML, NLVPMVATV; the immunodominant influenza-M1, EBV-BMLF1 and CMV-pp65 epitopes) are also among the VDJdb working labels, so the two TCR databases are not fully independent. The underlying TCR records differ (distinct curation sources and CDR3 sets), and the alignment-over-PLM ordering holds separately on each database; we therefore treat them as two partially overlapping replications rather than as wholly independent evidence.

#### 1.2 PLM embedding pipeline

Embeddings were precomputed in a single pass per model and stored as fixed-shape NumPy arrays. For each model, the final hidden state at the last transformer block was extracted. Mean pooling averaged over residue token positions, excluding the special beginning-of-sequence (BOS), end-of-sequence (EOS), and padding tokens. CLS pooling used the CLS/BOS token embedding directly. Two region-restricted poolings averaged over a subset of residue positions: CDR-only mean pooling over the CDR1/CDR2/CDR3 token positions, and framework-only mean pooling over the variable-domain non-CDR (framework) token positions. Region indices were determined by ANARCI (Dunbar and Deane, 2016) IMGT (Lefranc *et al.*, 2009) alignment for BCR/antibody inputs (CDRs at IMGT positions 27–38, 56–65 and 105–117; framework elsewhere within the numbered domain) and by IMGT CDR3 flanking-residue detection for TCR inputs. Embedding dimensionalities: ESM2-150M = 640, ESM2-650M = 1280, ESM2-3B = 2560, AntiBERTy = 512, AbLang = 768, TCR-BERT = 768. TCR-BERT (`wukevin/tcr-bert`) uses a whitespace-splitting tokenizer that expects space-separated single amino acids; we therefore space-separate residues before tokenizing. This model has a 64-token positional limit and a 20-amino-acid vocabulary, so it is valid on CDR3 amino-acid input. Because CDR3 and the VJ- and V-clonotype variants all rank by the CDR3 (the gene enters VJ- and V-clonotype only as a filter, §2.3), TCR-BERT is reported at CDR3, VJ-clonotype and V-clonotype; the paratope (concatenated CDRs) fits within its 64-token limit and is reported as an out-of-distribution point (Table S15), whereas the spaced full-length input exceeds the limit and is not assessable.

Cosine similarity was computed between pooled embeddings; for retrieval, candidate sequences were ranked by descending cosine similarity to the query embedding. Per-method retrieval did not include any task-specific fine-tuning.

**SCEPTR (contrastive TCR language model).** SCEPTR (Nagano *et al.*, 2025) is a TCR-specific language model pre-trained with an autocontrastive objective; it encodes the V gene together with the CDR3 into a single 64-dimensional embedding, so it is a fixed V+CDR3 representation rather than a per-residue model with a tunable representation. Its published package could not be installed alongside our main embedding environment (incompatible `torch`/`numpy`/CUDA requirements), so SCEPTR was run in an isolated CPU environment purely to *generate* embeddings, which were saved as offline NumPy arrays and scored through the *identical* clone-aware pipeline (`run.py`/`split.py`/`evaluate.py`; same 20 seeds, same expected Recall@1) as every other method. We used the `sceptr.variant.default()` model for both TCR databases (VDJdb mixes beta-chain and alpha-only CDR3-level records, and only the default variant embed all of them in one 64-dimensional space; on McPAS the default and beta-specific variants agree to within 0.001 Recall@1, so the choice is immaterial). V- and J-gene names were standardized with `tidytcells` before embedding; VDJdb required essentially no imputation, whereas McPAS's coarser gene calls required imputing ≈ 3% of V genes (and <1% of J genes and CDR3s). Because SCEPTR is a fixed V+CDR3 representation, it is assessable only at the CDR3 (CDR3) and the two clonotype variants (VJ-clonotype, V-clonotype); it has no multi-CDR (paratope) or full-length (full variable domain) input. As a direct, ranking-side check of its germline dependence, across all VDJdb queries SCEPTR's top-ranked reference shared the query's TRBV gene family 76% of the time, versus 40% for BLOSUM62 and 8% expected by chance (computed over every query, irrespective of whether the retrieval was correct, and therefore distinct from the correct-hit enrichment reported in §3.5). SCEPTR thus ranks by the encoded germline V gene directly, not only when it succeeds, which is why its CDR3-level edge over CDR3-only alignment is germline V context and disappears under the clonotype filter.

**Common gene-mappable comparison set (§3.5).** To compare SCEPTR against the purpose-built `tcrdist3` on exactly equal footing, the head-to-head reported in §3.5 scored all five methods (BLOSUM62, Levenshtein, ESM2-150M, SCEPTR and TCRdist) on the same per-seed clone-aware split *and* restricted every method to the common subset of test queries whose V/J genes `tcrdist3` can map (`tcrdist3` silently drops gene-unmappable records: 198/199 McPAS, 253/272 VDJdb single-chain, 113/199 VDJdb paired). Because this gene-mappable subset is modestly easier or harder than the full test set for individual methods, the §3.5 Recall@1 values differ slightly from the canonical full-set values in Table S1 (for example, VDJdb single-chain BLOSUM62 is 0.311 on the common subset versus 0.375 on the full set). Within the common subset the ordering is TCRdist ≈ SCEPTR > BLOSUM62 > Levenshtein > ESM2-150M at single-chain CDR3 on both databases, and TCRdist > SCEPTR ≈ BLOSUM62 on paired chains (full numbers in Table S14). SCEPTR is thus the only language model in our benchmark to reach alignment-level CDR3 retrieval, matching the germline-informed `tcrdist3` but not exceeding it; the edge disappears once the same germline V-gene context is given to alignment through the clonotype filter (VJ- and V-clonotype).

#### 1.3 Alignment and distance scores

**Levenshtein distance.** Computed with `rapidfuzz` (`rapidfuzz.distance.Levenshtein`), the standard unit-cost edit distance (insertion = deletion = substitution = 1; identical results to `python-Levenshtein`). Normalized by the length of the longer of the two sequences; rankings use ascending normalized distance.

**BLOSUM62 Smith–Waterman.** Local alignment with the BLOSUM62 substitution matrix, gap open penalty = 10 and gap extension penalty = 1, computed with `parasail` (Daily, 2016) (`parasail.sw_striped_16`). Two normalizations are reported: max-length normalization (used in the canonical analyses) and Karlin–Altschul bit-score (length-independent; used in sensitivity analyses reported in the released supplementary tables).

**TCRdist.** The paired-chain TCRdist baseline (Table S5) was computed with the canonical `tcrdist3` package (`TCRrep`): the CDR1, CDR2 and CDR2.5 loops are inferred from each chain's assigned V-gene and combined with the CDR3 under the standard IMGT position weights, and the paired distance is the sum of the alpha- and beta-chain TCRdist (the single-chain baseline uses the beta chain only). Retrieval ranks candidates by ascending TCRdist.

**BCRdist (a borrowed TCR metric, *not* a BCR-tuned method).** TCRdist (Dash *et al.*, 2017) and its `tcrdist3` implementation are designed and parameterized specifically for *T-cell* receptors (the CDR2.5 loop, the per-CDR position weights and the germline-gene handling are all TCR-specific), and the `tcrdist3` `TCRrep` class does not accept antibody (immunoglobulin) genes; `pwseqdist`'s Numba `tcrdist` kernel also failed to load in our environment. We therefore only *borrowed* the tcrdist single-CDR kernel (a faithful re-implementation: mid-gap alignment, ntrim/ctrim trimming, tcrdist substitution costs, CDR3 weighted 3×) and applied it to the ANARCI-extracted antibody CDR1/CDR2/CDR3, on the *identical* canonical clone-aware slices used by every other method, purely as a rough reference. On the same test queries this borrowed metric did not exceed the alignment baselines (BCR IEDB 0.499 vs. BLOSUM62 0.532 / Levenshtein 0.542; SAbDab single-chain 0.448 vs. 0.452; SAbDab paired 0.447 vs. 0.449). *Because tcrdist is not tuned for antibodies, we draw no conclusion about how a purpose-built BCR distance would perform*; we report this only to show that naively transplanting the TCR metric onto antibodies does not beat simple alignment. A genuine antibody-specific “BCRdist” analogous to `tcrdist3` does not exist in the literature and is left to future work.

**NCBI BLAST+ (convergent baseline).** As a second alignment baseline independent of BLOSUM62 Smith–Waterman, we used NCBI BLAST+ 2.16.0 (Altschul *et al.*, 1990) (`blastp`) with the `blastp-short` task, which is optimized for queries ≤ 30 amino acids. Per seed, the clone-aware training partition was written to FASTA and indexed with `makeblastdb -dbtype prot`; queries were the test partition. Parameters: `-task blastp-short -evalue 1000 -word_size 2 -max_target_seqs 25 -comp_based_stats F`. Retrieval used the top-1 bit-score hit; queries with no hits were assigned rank ∞ and counted as failures. BLAST was evaluated at CDR3 only on all four datasets to provide a second alignment baseline at CDR3 representation (full results in Table S10).

#### 1.4 Statistical methods

**Nested bootstrap confidence intervals.** For each method-dataset-representation combination, R@1 was computed per seed, and 1,000 nested bootstrap replicates were drawn over (i) the seed pool and (ii) the per-seed query pool. The reported 95% CI is the 2.5–97.5 percentile of the nested bootstrap distribution of the mean. The query-resampling step follows the out-of-sample-prediction bootstrap of Tsamardinos *et al.* (2018); the outer seed-resampling level is an additional hierarchical step.

**Hedges’ *g* effect size.** Computed from per-seed R@1 means and standard deviations using the standard correction factor for small samples ($g = J(df) \cdot d$, where d is Cohen’s d and $J(df) = 1 - 3 / (4\,df - 1)$). Effect-size categories follow Cohen’s conventional thresholds: negligible < 0.20, small < 0.50, medium < 0.80, large ≥ 0.80.

**Two one-sided tests (TOST).** For BCR full variable domain, per-seed differences $d_i$ = R@1(ESM2-150M)$_i$ - R@1(BLOSUM62)$_i$ were computed for i = 1, ..., 20. The two one-sided t-tests assess (i) $H_{01}$: d_mean ≤ -Δ versus $H_{11}$: d_mean > -Δ, and (ii) $H_{02}$: d_mean ≥ +Δ versus $H_{12}$: d_mean < +Δ, with Δ = 0.05 R@1. Equivalence is established if both null hypotheses are rejected at α = 0.05.

**Statistical power.** The per-seed standard deviation of the BLOSUM62-ESM2-150M R@1 difference across the 20 canonical seeds is approximately 0.073 (BCR), 0.041 (VDJdb TCR), 0.083 (SAbDab) and 0.030 (McPAS-TCR). Under a two-sided paired t-test at α = 0.05 with n_seed = 20, the minimum detectable effect at 80% power is approximately ± 5 pp R@1 for BCR, ± 3 pp for VDJdb TCR, ± 5.5 pp for SAbDab (which has the smallest working set, n_test ≈ 59, and the most variable per-seed differences) and ± 2 pp for McPAS-TCR. Differences smaller than these thresholds should be interpreted as not statistically distinguishable in our design; the principal BLOSUM62-vs-ESM2-150M CDR3 gaps (BCR +10.2 pp, TCR +9.4 pp, SAbDab +7.2 pp, McPAS +11.5 pp) are well above their respective detection floors.

**Clone-definition sensitivity.** Adding V-gene identity to the CDR3-only clone definition increased the cluster count by only 2.8%, confirming that CDR3 similarity alone is a sufficient proxy for clonal relatedness. Reported R@1 point estimates are query-weighted (pooled over all test queries through the nested bootstrap), whereas Hedges' g is computed from per-seed R@1; for SAbDab, whose per-seed test sets vary widely in size, these two aggregations differ, although both yield the same alignment advantage over ESM2.

**Secondary metrics.** Beyond Recall@1 we also computed Recall@5 (the correct label appears among the top five retrievals) and the mean reciprocal rank (MRR). Both are reported per database at CDR3 and full variable domain in Supplementary Table S13: the method ranking is unchanged from Recall@1: at the CDR3 input (CDR3) BLOSUM62 leads on both secondary metrics on all four databases, and at full length (full variable domain) the methods converge into a narrow band, so Recall@1 is not what drives the comparison.

**Tie handling.** When several reference sequences share the top similarity score, we report the *expected* Recall@1 under uniform-random tie resolution (the query receives fractional credit equal to the proportion of tied candidates sharing its label). Such ties are common at CDR3, arising from duplicate or near-identical CDR3 sequences in the bank (integer alignment scores tie outright; identical CDR3s give identical PLM embeddings): a top-score tie affects 42% of BCR queries for BLOSUM62, 56% for Levenshtein and 36% for ESM2-150M, and 17%/7% of VDJdb TCR queries (BLOSUM62/ESM2-150M; median tie size 2). With ties this prevalent the expected (fractional) treatment is a substantive modeling choice rather than a technicality: order-dependent 0/1 scoring would make per-seed CDR3 estimates depend on arbitrary tie-breaking. Optimistic (any tied match counts) and pessimistic (all must match) tie-handling nonetheless produce nearly identical rankings; the expected tie-handling is the primary metric throughout.

***Box S1. Recommended reporting standards for immune-receptor retrieval benchmarks***

1. **Split clone-aware.** Cluster clonally related receptors (e.g. by CDR3 similarity) and assign whole clusters to one side of the split. Random splits inflate Recall@1 by 15–28 pp (main paper §3.1).

2. **Use an order-independent, tie-aware metric.** Report expected Recall@1 (or state the tie convention explicitly); top-score ties are common at the CDR3 level for every method (main paper §2.1).

3. **Always report the zero-shot alignment baseline.** BLOSUM62 / Levenshtein, with a BLAST control, are cheap and strong; PLM gains should be measured against them, not against a no-baseline absolute (main paper §3.2).

4. **Stratify by representation and germline.** Separate CDR3 from full-chain inputs and add per-antigen and germline- / SHM-stratified breakdowns. Aggregate full-length parity can hide a germline shortcut and per-antigen divergence (main paper §3.3–3.4).

5. **Sweep the clone threshold.** Report sensitivity to the clustering cutoff rather than relying on a single value (main paper §3.5).

6. **Report embedding provenance.** State the model version, the layer read, the pooling, and the tokenization; these choices change the numbers (a tokenization fix alone shifted one antibody PLM's CDR3 Recall@1 by ~14 pp).

### 2 Supplementary Notes

This section reports six supplementary analyses bearing on the germline-shortcut reading, cross-referenced from the main paper Results (§3). They group by the claim they support: **S2.1** and **S2.6** concern the CDR3-level deficit (its scale-dependence; and that it is representational, not a metric or geometry artifact); **S2.2**, **S2.4** and **S2.5** are the three independent controls behind the full-length germline shortcut (the framework×representation interaction with pooling robustness; the V-gene oracle; and the germline-reversion test); and **S2.3** examines the HIV-1/somatic-hypermutation exception. Notes are numbered as S2.1–S2.6 in the auto-generated section numbering and are also referred to in the running text as Note S2.1, etc.

#### 2.1 McPAS-TCR scale-dependent retrieval

On McPAS-TCR CDR3, all three ESM2 scales remained well above the 10-label random-retrieval baseline (R@1 = 0.100, the sum of squared test-label frequencies, equal to 1/10 for the near-balanced capped label set) and well below the BLOSUM62 reference (0.387). In contrast to BCR and SAbDab (where increasing ESM2 size produced no consistent CDR3 improvement), McPAS-TCR showed a modest monotonic gain with scale: R@1 = 0.272 at 150M, 0.282 at 650M and 0.305 at 3B (**Supplementary Figure** S6). The ESM2 values sit far above this random floor, confirming that the absolute scale is meaningful rather than a data-processing artifact, yet the largest model still trails alignment by 8.2 pp.

This dataset-specific scaling behavior indicates that the effect of model size on zero-shot CDR3 retrieval depends on the receptor and antigen context rather than following a single scaling law. Even at the largest scale, however, ESM2 did not approach the alignment baseline on McPAS-TCR, consistent with the main-text finding that increasing model size alone does not close the CDR3-level gap between PLM embeddings and alignment scoring.

#### 2.2 Pooling sensitivity and the framework–representation interaction

**Pooling sensitivity.** Four pooling strategies were compared on BCR full variable domain using ESM2-150M, on the cleaned variable-domain inputs (20-seed canonical splits, expected-R@1):

1. **Mean pooling** (canonical): average over all residue tokens. R@1 = 0.507 [95% CI 0.471–0.544].
2. **CLS pooling**: use the CLS token embedding. R@1 = 0.484 [95% CI 0.452–0.516].
3. **CDR-only mean pooling**: average over the CDR1/CDR2/CDR3 token positions only. R@1 = 0.529 [95% CI 0.490–0.568].
4. **Framework-only mean pooling**: average over the variable-domain non-CDR token positions only. R@1 = 0.489 [95% CI 0.447–0.531].

All four strategies were statistically indistinguishable (every pair of 95% confidence intervals overlaps) and CLS pooling showed no collapse (per-seed CLS minus mean = -2.4 pp), so the full variable domain parity result is robust to how embeddings are aggregated. (CLS pooling is sensitive to length heterogeneity: on the variable-domain-trimmed inputs used throughout it matches mean pooling and shows no collapse, whereas the untrimmed, Fab-contaminated sequences would make it unreliable. The sequences are trimmed to the variable domain for the antigen-correlated constant-region confound described in the Supplementary Methods, not specifically for CLS.) This test does *not*, however, localize the full variable domain signal to either region: under global self-attention every retained token has already attended to the rest of the chain, so neither CDR-only nor framework-only pooling removes the other region's information. The only directional difference is that CDR-only pooling was, if anything, marginally *higher* than framework-only (per-seed paired difference +4.0 pp, 95% CI [+1.4, +6.5]), the opposite of what a framework-localized signal would predict. We therefore treat pooling as a robustness / no-collapse check, not as evidence for where the signal resides. See **Supplementary Figure S2**.

**Framework–representation interaction.** The germline-shortcut reading rests instead on a direct input-ablation result: adding the germline framework (paratope, all CDRs → full variable domain, full chain) benefits the PLM but not alignment. Using per-seed R@1 (20 seeds, identical clone split, paired across representations and methods), the interaction (the paratope→full-variable-domain gain for the PLM minus that for alignment) is positive in every one of the 24 paired comparisons (four databases × three ESM2 sizes × two alignment baselines), each individually significant at $p<0.05$ (paired Hedges g 0.60–1.95). These 24 cells are not independent (within a database the comparisons reuse the same per-method Δ), so we do not combine them into a single p-value; the unit of independent replication is the four databases, in each of which every ESM2 size exceeds every alignment baseline. Twelve of the 24 comparisons are on the native BCR and SAbDab datasets (non-circular); the other twelve are on germline-reconstructed TCR and are consistent but not independent. Per-method paratope→full variable domain change (mean ΔR@1 over 20 seeds):

| Dataset | Levenshtein | BLOSUM62 | ESM2-150M | ESM2-650M | ESM2-3B |
|---|---|---|---|---|---|
| BCR (native) | -0.044 | -0.019 | +0.070 | +0.120 | +0.082 |
| SAbDab (native) | -0.033 | -0.027 | +0.043 | +0.021 | +0.054 |
| VDJdb (recon.) | -0.015 | -0.009 | +0.022 | +0.034 | +0.021 |
| McPAS (recon.) | -0.037 | -0.050 | +0.037 | +0.007 | +0.003 |

Alignment never gained from the framework (8/8 datasets×methods with Δ≤ 0, significantly negative on McPAS and SAbDab); the PLM gained throughout. The two TCR datasets (VDJdb, McPAS) have germline-reconstructed full variable domain, so their added framework is synthetic; the native datasets (BCR, SAbDab) show the same interaction. Two database-dependent nuances are consistent with the PLM's gains coming from germline regions that plateau once captured: in SAbDab the germline CDR1/CDR2 add no retrieval signal (CDR3→paratope ≈ 0 for all methods) because its CDR3s are the most within-antigen-redundant of the four datasets (a few dense structural-antibody families), so CDR3 retrieval already saturates, not because the V-gene is uninformative there (SAbDab has the strongest V-gene–antigen association, Cramér's V = 0.69); and in McPAS, where CDR1/CDR2 already lift the PLM strongly, the larger ESM2 models gain little further from the framework.

#### 2.3 SHM-stratified BCR full variable domain analysis

Somatic hypermutation (SHM) density was computed per BCR full variable domain test sequence as the fraction of mismatches in IMGT V-region positions 1–104 relative to the ANARCI-assigned germline allele. SHM density was binned as Low (<5%), Medium (5–15%) and High (≥ 15%). Per-bin R@1 was computed from 20 canonical seeds; see Table S9.

The crossover at the High SHM bin (ESM2-150M -9.0 pp at Low, -1.8 pp at Medium, but +7.0 pp at High) is the largest divergence observed in the BCR full variable domain evaluation. Two interpretive caveats apply:

- **Antigen confounding, addressed by within-antigen stratification**: the High SHM bin is substantially composed of HIV-1 envelope sequences (estimated ~ 63% from per-antigen test-set sizes), and HIV-1 is precisely the antigen on which ESM2-150M shows its largest per-antigen advantage (§3.4). To separate the two, we stratified by SHM *within* a single antigen (pooling query instances across the 20 seeds). Inside HIV-1 envelope the ESM2-minus-BLOSUM62 R@1 advantage is clearly positive at medium and high SHM (Low -0.026, Medium +0.386, High +0.186; n = 39/153/363 pooled instances; +0.226 overall), though not strictly monotonic. This pattern does *not* generalize, however: inside influenza hemagglutinin the advantage is slightly negative (Low -0.070, High -0.059), and restricting to the High-SHM bin the ESM2 advantage is absent for non-HIV antigens (-0.028, n = 419). The high-SHM crossover therefore appears to be largely an HIV-1-envelope effect (where high SHM density and ESM2's per-antigen advantage coincide) rather than a general SHM-density rule; with these data we cannot cleanly separate an SHM-density contribution from HIV-1 antigen identity. These strata pool non-independent query instances across seeds and are not formally powered per seed, so we report direction only.

- **Pretraining leakage**: HIV-1 broadly neutralizing antibody (bNAb) CDR3 sequences (such as VRC01 and VRC03) are present in UniRef90 and may have been seen by ESM2 during pretraining. A leave-one-out experiment excluding all CDR3 sequences with ≥ 80% Levenshtein identity to a panel of known bNAbs (7 unique sequences, appearing in 8 of the 20 seeds) yields a difference-in-differences memorization effect = +1.1 pp (removing the flagged sequences slightly *increases*, rather than decreases, ESM2's relative advantage), indicating no detectable memorization-specific advantage. The +22.2 pp leak-free ESM2 advantage on HIV-1 envelope (0.634 vs. 0.412 on the cleaned subset) is therefore authentic.

With the within-antigen stratification above, the high-SHM crossover is best read as largely an HIV-1-envelope effect: the ESM2 advantage is strong within HIV-1 at medium and high SHM but does not reproduce inside influenza or in non-HIV high-SHM sequences. We therefore attribute it to the HIV-1-antigen×SHM regime rather than a general SHM-density rule, and stop short of a calibrated causal claim because the within-antigen strata pool query instances across seeds and are not powered for per-seed inference.

#### 2.4 V-gene confusion and germline shortcut

At full-length (full variable domain), 91.5% (ESM2-150M) to 99.2% (Levenshtein) of top-1 retrievals shared the query's V-gene family, versus ~ 34.0% expected at random; at CDR3-only representation (CDR3) this fell to 53.7%–63.2% (Table S11). The increase at full variable domain reflects the addition of germline-encoded framework residues, which strongly bias retrieval toward same-V candidates.

A V-gene-only analytical baseline computed as the expected R@1 under uniform random selection from same-V-gene-family training candidates gives R@1 = 0.190 [95% CI 0.181–0.200]. Sequence-based methods (0.498–0.513 on this subset) substantially exceed this baseline by ~ 0.32 R@1 units, indicating that within-germline sequence discrimination contributes genuine additional signal.

A two-stage oracle (Stage 1: filter by V-gene family; Stage 2: rank by full-length BLOSUM62) achieves R@1 = 0.507 [95% CI 0.464–0.550], matching the benchmark methods. Critically, this does *not* exceed full variable domain BLOSUM62 (0.522 on the full variable domain set; 0.513 on the same V-gene-assignable subset used here); if anything it is marginally lower, despite the oracle using explicit V-gene ground truth. This decomposition indicates that full-length BLOSUM62 already implicitly captures the V-gene-identity signal, consistent with the full variable domain convergence between ESM2-150M and BLOSUM62 arising from both methods exploiting the same germline signal rather than from PLM-learned CDR-level discrimination.

#### 2.5 Germline-reversion test

To test directly whether the PLM's full-length gain is germline-driven, we reverted one region of each V-gene-assignable antibody to its assigned germline IGHV allele (IMGT-gapped reference `IGV.fasta`) while keeping the rest native, and re-evaluated under the canonical 20-seed clone-aware protocol (expected-R@1). Two variants were built: **germ-fw** (germline FW1–FW3 + native CDR1/CDR2/CDR3 + native FW4; framework reverted) and **germ-cdr** (native framework + germline CDR1/CDR2 + native CDR3; CDR1/2 reverted). All inputs were embedded on identical splits; absolute R@1 on this V-assignable subset runs slightly below the canonical full set, so we report per-seed paired *differences*, which are offset-independent.

**Decomposition (ESM2-150M, BCR, 20-seed paired ΔR@1).** paratope → germ-fw (germline framework) +0.069 [+0.048,+0.090], $p<10^{-5}$; germ-fw → full variable domain (framework somatic mutation) +0.039 [+0.006,+0.073], p=0.034; germ-cdr → full variable domain (CDR1/2 somatic mutation) +0.023 [-0.008,+0.053], ns; total paratope → full variable domain +0.108, $p<10^{-5}$. For BLOSUM62 every step was within noise of zero. The germline framework thus accounts for about two-thirds of the PLM's framework gain, while somatic mutation in CDR1/CDR2 contributes nothing (Figure S9).

**Antigen stratification and leakage.** The framework-somatic gain is entirely HIV-1 envelope: +0.183 [+0.110,+0.256] for HIV queries versus -0.012 (ns) for all other antigens. After removing the four bNAb-matched CDR3s present in this subset (among the seven flagged at ≥ 80% identity in Note S2.3), the HIV effect falls to +0.113 [+0.033,+0.193], p=0.012 (about 38% of the raw effect was pretraining memorization) but remains significant; the germline-framework gain is unchanged (+0.076). The PLM therefore reads genuine somatic-framework signal only in the most hypermutated antibodies.

**Cross-database replication (SAbDab).** The germline-framework gain replicated on the native SAbDab antibodies (paratope → germ-fw +0.050 [+0.005,+0.094], p=0.04). There, however, framework somatic mutations *reduced* ESM2 retrieval (germ-fw → full variable domain -0.070, p=0.002): SAbDab's structure-derived, engineered/humanized frameworks diverge from germline in non-natural ways that the PLM mis-reads. The framework-somatic "real signal" is thus HIV-specific and does not generalize.

**Length-matched framework-CDR3 control (framework-CDR3 control).** As an independent control we replaced CDR1/CDR2 with matched-length glycine spacers while retaining FW1–FW4 and the native CDR3. For ESM2-150M, paratope (0.445) → framework-CDR3 control (0.474; +2.9 pp) → full variable domain (0.514; +4.0 pp); for BLOSUM62 the framework added nothing (0.538 → 0.517 → 0.522), so alignment already resolves antigen identity from the CDRs at paratope. Because the paratope → framework-CDR3 control step changes two things at once (adds framework, replaces CDR1/2 with out-of-distribution glycine) it is less clean than the germline-reversion above, but points the same way (Figure S8, Table S3).

**Validation.** germ-fw and germ-cdr preserve the native CDR3 (100%), parse as valid variable domains (ANARCI), and match native length. The within-V-gene collapse from germline-framework reversion does not create train/test leakage: the germ-fw test→train exact-duplicate rate (18.1%) equals that of CDR3/paratope/full variable domain (17.7–19.3%), and the effect is PLM-specific (BLOSUM62 flat at every step), ruling out a trivial "cleaner-sequence" artifact. Scripts: `build_germline_framework_bcr.py`, `run_germfw_experiment.py`, `run_germfw_sabdab.py`, `verify_germfw.py`.

**Germline-subtracted residual.** As a complementary localization of the somatic signal, we subtracted the germline-reverted (germ-fw) embedding from the native full variable domain embedding to isolate the framework's somatic component, and retrieved on this residual alone. The residual retrieves HIV-1 envelope at R@1 =0.596 [0.512, 0.680], close to full native full variable domain (0.575), but other antigens at only 0.299 [0.264, 0.335] (random 0.100). The somatic-framework signal is thus about twice as informative for the hypermutated HIV-1 antibodies as for the rest, sharpening the germline-reversion result above. These residual numbers use the full HIV-1 query set (no leak exclusion); because the leak-excluded germline-reversion effect above remains significant at p=0.012, the direction is robust, though the absolute residual R@1 may be slightly inflated by the ~38% memorization component identified in the raw framework-somatic step (`scratch/germline_residual_experiment.py`).

#### 2.6 CDR3-level deficit: pseudo-perplexity and germline-distance tests

We tested whether the zero-shot PLM's CDR3 deficit reflects the non-evolutionary, junctional nature of the CDR3. For each BCR CDR3 query we computed ESM2-150M's masked-language pseudo-perplexity (mask each residue in turn, PPL=exp(-(1/L)$\Sigma_i$ log P($x_i$ | x_≠ i))), a per-sequence measure of how out-of-distribution the CDR3 is to the model, alongside the per-query expected-R@1 for ESM2-150M and BLOSUM62 (20 seeds, 2,564 query instances, 559 unique CDR3s; `scratch/ood_cdr3_experiment.py`).

The PLM's per-query advantage over BLOSUM62 falls as pseudo-perplexity rises. Because the 2,564 per-query instances are 20-seed resamples of 559 unique CDR3s and are not independent, we report the association at the level of the 559 unique CDR3s (mean over seeds): Spearman ρ=-0.14, p=$6\times10^{-4}$. The seed-pooled per-query value is more extreme (ρ=-0.18, Pearson -0.16) but is inflated by this non-independence and quoted only descriptively. This is not a CDR3-length artifact: pseudo-perplexity is only weakly related to length (ρ=-0.25), the deficit is uncorrelated with length (ρ=+0.00), and the partial correlation of the deficit with pseudo-perplexity controlling for length is unchanged (-0.185). By pseudo-perplexity tertile (Figure S10):

| Tertile | mean PPL | ESM2-150M R@1 | BLOSUM62 R@1 | deficit (ESM2-BLOSUM) |
|---|---|---|---|---|
| low (in-distribution) | 17.9 | 0.490 | 0.513 | -0.023 |
| medium | 20.5 | 0.319 | 0.377 | -0.058 |

| Tertile | mean PPL | ESM2-150M R@1 | BLOSUM62 R@1 | deficit (ESM2-BLOSUM) |
|---|---|---|---|---|
| high (out-of-distribution) | 23.8 | 0.261 | 0.489 | -0.229 |

ESM2-150M retrieval declines monotonically with pseudo-perplexity while BLOSUM62 does not decline monotonically (it dips in the medium tertile and recovers in the high tertile); the two are at parity on the most natural-like CDR3s and diverge by 23 pp on the ones ESM2 scores as least typical. Whether this model-assessed atypicality reflects the junctional, non-germline diversity that V(D)J recombination adds to the CDR3 (random N/D-region additions), as one might expect, is tested directly below.

**Model-independent check.** Because pseudo-perplexity is internal to ESM2, we tested a biological axis that does not rely on the model: each CDR3's *germline-distance*, its normalized edit distance to the nearest germline V–D–J recombination (`scratch/ood_germline_distance.py`). D-gene assignment and N1/N2 junction segmentation are notoriously unreliable, so we do not assign a D or parse the junction: we take the minimum over all IMGT IGHD genes (and the no-D V–J join) and let the edit distance absorb D-end trimming (as deletions) and N/P additions (as insertions); the residual distance is then the CDR3 content not explained by *any* germline V–(D)–J join (IGHD is available only as its frame-1 translation, one further limitation). Including germline D removes most of the length dependence of this axis (Spearman with length +0.14, versus +0.39 for a V–J-only join). Even so, germline-distance does *not* track the deficit (Spearman -0.03, p=0.4 over the 559 unique CDR3s; -0.04 partialling out length; robust to using only functional D genes) and is itself *uncorrelated* with pseudo-perplexity (+0.03, p=0.5). With n=559 powered to detect the pseudo-perplexity effect (ρ=-0.14), this is a genuine null rather than low power. We therefore keep only the model-internal statement: the deficit concentrates on CDR3s ESM2 scores as atypical, but this atypicality does *not* correspond to a model-independent germline or junctional distance, so we do not attribute the deficit to junctional germline-divergence. A stronger biological test (all three D reading frames; a per-residue sequence-naturalness score) remains future work.

**Geometry versus representation.** We next asked whether the CDR3 deficit is a property of the cosine *metric* on anisotropic embeddings (a fixable geometry artifact) or of the *representation* itself (`scratch/cdr3_mechanism_geometry.py`, `cdr3_mechanism_supp.py`; same canonical pipeline and 20 seeds, so the raw baselines reproduce the main values; all transforms fit on the reference set only). Zero-shot CDR3 embeddings are extremely anisotropic (mean pairwise cosine 0.95–0.99 across ESM2-150M/650M/3B and ESM-C; the top principal component alone is ~24% of variance), so cosine geometry is a natural suspect. It is not the cause. Standard isotropy restoration, whether centering, removing the top principal components ("all-but-the-top"), or PCA-whitening in the top-r reference subspace, recovers little: centering and top-PC removal are within noise, and PCA-whitening, the only significant fix, adds just +0.03 to +0.05 Recall@1 and leaves every model well below alignment (table below; Figure S11). Two further controls rule geometry out directly: (i) the full-chain (full variable domain) embeddings, where the PLM is at *parity* with alignment, are just as anisotropic (0.97–0.99) yet show no deficit and no whitening gain; and (ii) the CDR3 deficit persists under Euclidean distance (ESM2-150M raw 0.360 versus cosine 0.353). The deficit is therefore representational: the zero-shot embedding under-encodes the discriminative junctional signal, and it is not an artifact of comparing anisotropic vectors by cosine. This is the direct, geometry-controlled version of the linear-probe observation (main text §3.2).

| Model (BCR CDR3) | anisotropy | raw R@1 | PCA-whitened R@1 | gap to BLOSUM62 (0.459) |
|---|---|---|---|---|
| ESM2-150M | 0.95 | 0.353 | 0.384 (+0.031, t=2.9) | -0.075 |
| ESM2-650M | 0.98 | 0.369 | 0.418 (+0.049, t=4.0) | -0.041 |
| ESM2-3B | 0.99 | 0.344 | 0.369 (+0.025, t=2.5) | -0.090 |
| ESM-C-300m | 0.96 | 0.374 | 0.381 (+0.008, n.s.) | -0.078 |
| ESM2-150M, full variable domain (parity) | 0.98 | 0.507 | 0.503 (n.s.) | — |

Anisotropy = mean pairwise cosine of reference embeddings; whitening (top-128 reference subspace) fit on the reference set, applied to both sides; t = paired across 20 seeds. Even the best fix leaves every CDR3 PLM below alignment; full variable domain is equally anisotropic yet at parity.

### 3 Supplementary Tables

This section reports fifteen supplementary tables. Table S1 is the master results table (expected Recall@1 for every dataset × representation × method, all four databases plus paired chains). Tables S2–S11 provide dataset composition (S2), the BCR representation ladder including the framework-CDR3 control (S3), per-seed TOST equivalence test (S4), paired-chain results (S5), threshold sensitivity (S6), per-antigen BCR CDR3 (S7) and full variable domain (S8), SHM-stratified BCR full variable domain (S9), BLAST CDR3 cross-reference (S10), and V-gene sharing rates (S11); Table S12 summarizes, for every benchmark representation, what it isolates and what its result indicates; Table S13 reports the secondary retrieval metrics (Recall@5 and MRR); Table S14 gives the SCEPTR vs. alignment vs. `tcrdist3` head-to-head on the common gene-mappable subset (§3.5); and Table S15 lists the domain-specific and auxiliary models (companion to S1).

***Table S1. Master retrieval results: expected Recall@1 across all datasets and representations.*** *Twenty-seed bootstrap means of expected Recall@1 (order-independent tie handling, §2.1) for the five benchmark methods assessable at every* *representation. Representations follow the defined order (§2.3): the input representations CDR3, paratope (all CDRs), framework-CDR3 control (BCR only), and full variable domain (full chain), interleaved with the two clonotype-retrieval variants VJ-clonotype (V+J germline-gene filter) and V-clonotype (V-gene filter). "Paired" rows use both receptor chains. Within each dataset the alignment methods (Levenshtein, BLOSUM62) lead at the CDR3 input and peak by the paratope, while the PLM rises toward full variable domain: the BCR framework-CDR3 control indicates this rise draws on both the germline framework and the germline-derived CDR1/CDR2 loops (§3.3). The same ladder pattern holds across VDJdb TCR, McPAS-TCR and SAbDab; framework-CDR3 control is BCR-only (see footnote). SAbDab paired now reports the full ladder (CDR3 to the full variable domain) (antibody heavy+light CDR3 and all-CDR inputs; paired VJ- and V-clonotype filter uses both heavy and light V/J genes). Domain-specific models, the purpose-built TCRdist3, and a BCRdist negative control are summarized in the footnote; full 95% confidence intervals for every cell are in the released `Table_S1_Full_CI.csv`. "—" marks a representation that is not validly assessable (see footnote and Supplementary Methods).*

| Dataset | Level | Levenshtein | BLOSUM62 | ESM2-150M | ESM2-650M | ESM2-3B |
|---|---|---|---|---|---|---|
| BCR (IEDB) | CDR3 | 0.446 | 0.459 | 0.357 | 0.372 | 0.350 |
| | VJ-clonotype | 0.432 | 0.433 | 0.397 | 0.408 | 0.412 |
| | V-clonotype | 0.517 | 0.526 | 0.483 | 0.482 | 0.466 |
| | paratope | 0.550 | 0.538 | 0.445 | 0.417 | 0.446 |
| | framework-CDR3 control‡ | 0.509 | 0.517 | 0.474 | 0.514 | 0.487 |
| | full variable domain | 0.508 | 0.522 | 0.514 | 0.541 | 0.528 |
| VDJdb TCR (single) | CDR3 | 0.355 | 0.375 | 0.282 | 0.304 | 0.307 |
| | VJ-clonotype | 0.310 | 0.315 | 0.295 | 0.305 | 0.300 |
| | V-clonotype | 0.347 | 0.356 | 0.323 | 0.326 | 0.334 |
| | paratope | 0.387 | 0.406 | 0.352 | 0.371 | 0.367 |
| | full variable domain | 0.384 | 0.400 | 0.390 | 0.393 | 0.392 |
| VDJdb TCR (paired) | CDR3 | 0.414 | 0.436 | 0.352 | 0.349 | 0.363 |
| | VJ-clonotype | 0.123 | 0.123 | 0.123 | 0.122 | 0.123 |
| | V-clonotype | 0.180 | 0.179 | 0.178 | 0.178 | 0.180 |
| | paratope | 0.442 | 0.458 | 0.395 | 0.396 | 0.399 |
| | full variable domain | 0.437 | 0.437 | 0.424 | 0.427 | 0.430 |

| Dataset | Level | Levenshtein | BLOSUM62 | ESM2-150M | ESM2-650M | ESM2-3B |
|---|---|---|---|---|---|---|
| McPAS-TCR | CDR3 | 0.360 | 0.387 | 0.272 | 0.282 | 0.305 |
| | VJ-clonotype | 0.385 | 0.389 | 0.388 | 0.381 | 0.387 |
| | V-clonotype | 0.479 | 0.495 | 0.468 | 0.467 | 0.478 |
| | paratope | 0.461 | 0.473 | 0.392 | 0.409 | 0.407 |
| | full variable domain | 0.424 | 0.426 | 0.427 | 0.415 | 0.414 |
| SAbDab | CDR3 | 0.351 | 0.358 | 0.286 | 0.273 | 0.259 |
| | VJ-clonotype | 0.266 | 0.261 | 0.265 | 0.256 | 0.253 |
| | V-clonotype | 0.348 | 0.349 | 0.333 | 0.333 | 0.317 |
| | paratope | 0.363 | 0.378 | 0.292 | 0.285 | 0.295 |
| | full variable domain | 0.330 | 0.351 | 0.335 | 0.306 | 0.349 |
| SAbDab (paired) | CDR3 | 0.377 | 0.359 | 0.299 | 0.289 | 0.286 |
| | VJ-clonotype | 0.122 | 0.123 | 0.121 | 0.121 | 0.121 |
| | V-clonotype | 0.214 | 0.215 | 0.215 | 0.219 | 0.211 |
| | paratope | 0.379 | 0.367 | 0.280 | 0.303 | 0.269 |
| | full variable domain | 0.357 | 0.360 | 0.352 | 0.353 | 0.355 |

‡ **framework-CDR3 control (BCR only):** the length-matched framework-CDR3 control replaces CDR1 and CDR2 with glycine spacers of matched length while retaining FW1–FW4 and the native CDR3 (see Supplementary Note S2.5 and Supplementary Figure S8). framework-CDR3 control is not computed for VDJdb TCR, McPAS-TCR or SAbDab because those datasets use IMGT-reconstructed full-length sequences (VDJdb, McPAS) or structure-derived sequences (SAbDab) without separate CDR1/CDR2 loop annotations; this substitution control is therefore BCR-specific.
**TCR paratope (VDJdb, McPAS):** because the paratope input concatenates CDR1+CDR2+CDR3, which requires a V-gene call to obtain CDR1/CDR2, the TCR and McPAS paratope rows are computed on the subset of records with an assignable V gene; their values therefore differ by ≤1–2 pp from a naive recompute on the full set, while preserving the alignment-over-PLM ordering. BCR paratope uses the full set.
*Domain-specific and auxiliary models*. Per-model expected Recall@1 (20-seed nested-bootstrap means) for AbLang, AntiBERTy, TCR-BERT, SCEPTR and ESM-C-300m is given in Table S15. None beats alignment at the CDR3 except SCEPTR, and it does so only through its encoded germline V-gene context: it converges with or falls below alignment at the gene-filtered clonotype variants, and its head-to-head against `tcrdist3` on the common

gene-mappable subset is in §3.5 and Table S14. ESM-C-300m, a newer general-purpose PLM (EvolutionaryScale, 2024-12; the smallest ESM-C size; mean pooling, 20-seed nested), is statistically on par with ESM2-150M at the CDR3 (7.1–9.9 pp below BLOSUM62 on all four databases), converges with the other methods at full length, and shows no CLS-token collapse (CLS/mean R@1 ratio 0.97–1.08; the better CLS variant, BCR CDR3 0.419, still trails BLOSUM62 0.459), so this newer architecture does not change the conclusions. TCRdist3 (`tcrdist3`, real CDR-weighted distance): VDJdb single-chain (beta) full variable domain = 0.428; VDJdb paired (α+β) full variable domain = **0.515**, the highest paired value of any method tested. McPAS-TCR full variable domain = 0.253† (directional evidence only: 78.9% of rows retained after V-gene normalization; dominant-epitope clonal skew produces identical train/test partitions across all 20 seeds, so no nested-bootstrap CI is available). BCRdist (a *borrowed* TCR metric, not a BCR-tuned method; see Supplementary Methods): the tcrdist3 distance is TCR-specific and does not accept antibody genes, so we only borrowed its single-CDR kernel and applied it to antibody CDR1/CDR2/CDR3 on the identical canonical slices as a rough reference. This borrowed metric does *not* exceed simple alignment on antibodies (BCR IEDB 0.499 vs. BLOSUM62 0.532 / Levenshtein 0.542; SAbDab single 0.448 vs. 0.452; SAbDab paired 0.447 vs. 0.449). We draw no conclusion about a purpose-built BCR distance; it is reported only to show that transplanting the TCR metric onto antibodies does not beat alignment, and is not a benchmark row. SAbDab (paired): VJ-clonotype values are low (≈ 0.12) because the two-chain V+J germline-gene filter (VJ-clonotype) is extremely strict for paired antibody chains (requiring the query and candidate to share both heavy *and* light chain V and J genes), and most query–candidate pairs fail the filter (miss); the convergence across methods reflects the filter dominating the score, not the similarity metric. V-clonotype (V-gene only) partially relaxes this and recovers to ≈ 0.22. This mirrors the pattern observed in VDJdb TCR paired chains (main paper §3.3); at CDR3 and paratope the alignment methods lead the PLMs, as in every other dataset. *"—" (not assessable) entries, by reason*. **Receptor type:** AbLang and AntiBERTy are antibody-specific PLMs and are not applied to TCR datasets (VDJdb, McPAS); TCR-BERT is TCR-specific and is not applied to BCR or SAbDab. **Architecture/training regime:** TCR-BERT is reported at CDR3, VJ-clonotype, V-clonotype and, out of distribution for its CDR3-only training, the paratope (Table S15); only the full variable domain is not assessable, as the spaced full-length input exceeds its 64-token positional limit. **Method design:** TCRdist3 computes a single CDR1/CDR2/CDR2.5/CDR3-weighted distance with no input-representation ladder, so it is reported only at the full variable domain position where a direct comparison to the five main methods is meaningful; it is not reported at CDR3 to the paratope. **Data limitation (McPAS-TCR, partial):** McPAS-TCR V-gene annotations are heterogeneous in format (partial identifiers such as `TRBV4`, full identifiers such as `TRBV4-1*01`, and slash-ambiguous entries such as `TRBV2-1/TRBV2-2`). After normalizing the 78.9% of rows whose V genes can be resolved to a unique IMGT entry, TCRdist3 runs successfully and returns McPAS full variable domain R@1 = 0.253†. However, the dominant epitope (`lprrsgaaga`, 31% of rows) causes extreme clonal clustering: all 20 random seeds produce the *same* clone-aware train/test partition, so no cross-seed variance exists and no nested-bootstrap CI can be computed. This value is therefore reported as directional evidence only (footnote †), not as a full nested-bootstrap estimate. The 21.1% of rows excluded due to unresolvable V genes (ambiguous sub-gene family or not in the IMGT/tcrdist3 database) introduce a coverage caveat. Note: ANARCI supports both BCR and TCR variable-domain numbering and V/J germline assignment, but requires a full variable-domain sequence (FW1–CDR3–FW4); it cannot infer V genes from CDR3 sequence alone. McPAS provides only CDR3 sequences (no full variable domain), so ANARCI cannot be used to resolve the ambiguous V-gene annotations here. **Note on TCRdist heuristic:** an earlier CDR-weighted BLOSUM62 heuristic ("TCRdist surrogate") was used during development; it has been superseded by the real `tcrdist3` implementation and is not reported here. Full 95% CIs for all cells are in the released `Table_S1_Full_CI.csv`. Representation restrictions are detailed in Supplementary Methods.

***Table S2. Dataset composition after preprocessing.*** *Counts at each preprocessing stage. SAbDab raw count is heavy-chain structures in SAbDab; subsequent stages isolate clean paired human antibody–antigen records. McPAS-TCR is subsampled to the top 10 most-abundant epitope labels.*

| Dataset | Raw | Species filter | Label filter | Train (clone-aware) | Test (clone-aware) |
|---|---|---|---|---|---|
| BCR (IEDB) | 8,747 | 5,063 | 2,744 | ≈ 546 | ≈ 128 |
| TCR (VDJdb) | 139,744 | 126,137 | 92,600 | ≈ 801 | ≈ 199 |
| SAbDab | 23,466 | — | — | ≈ 251 | ≈ 59 |
| McPAS-TCR | 41,306 | — | 1,550 | ≈ 801 | ≈ 199 |

"—" marks a stage absent from that dataset's pipeline: SAbDab records are structure-derived and filtered directly to human paired antibody–antigen complexes, and McPAS-TCR is drawn from a pre-curated, species-annotated catalog. Train/Test columns are the per-seed clone-aware split sizes after all filtering.

***Table S3. BCR benchmark ladder in defined order: CDR3, VJ-clonotype, V-clonotype, paratope, framework-CDR3 control and full variable domain.*** *The input representations (CDR3, paratope, full variable domain) and the length-matched framework-CDR3 control are interleaved with the two clonotype-retrieval variants (VJ-clonotype, V+J; V-clonotype, V only) at their numerical positions. Recall@1 with 95% nested-bootstrap CI across 20 canonical seeds. framework-CDR3 control is the length-matched framework-CDR3 variant: CDR1 and CDR2 are replaced with glycine spacers of matched length, FW1–FW4 and the native CDR3 are retained. VJ- and V-clonotype are two-stage clonotype retrieval (see † and §2.3), not input representations. See Supplementary Figure S8 for visualization. Among the input representations, bold marks the highest value in each method column: the alignment methods peak at the all-CDR paratope, whereas the PLM peaks at full length (full variable domain).*

| Level | Input | Levenshtein R@1 | BLOSUM62 R@1 | ESM2-150M R@1 |
|---|---|---|---|---|
| CDR3 | CDR3 only | 0.446 [0.407, 0.487] | 0.459 [0.417, 0.498] | 0.357 [0.322, 0.393] |
| VJ-clonotype† | Clonotype: V+J filter → CDR3 | 0.432 [0.394, 0.473] | 0.433 [0.395, 0.474] | 0.397 [0.360, 0.435] |
| V-clonotype† | Clonotype: V filter → CDR3 | 0.517 [0.479, 0.554] | 0.526 [0.487, 0.562] | 0.483 [0.447, 0.522] |
| paratope | CDR1+CDR2+CDR3 | **0.550 [0.504, 0.589]** | 0.538 [0.503, 0.572] | 0.445 [0.401, 0.488] |
| framework-CDR3 control | Framework only (CDR1/CDR2 → Gly) | 0.509 [0.460, 0.555] | 0.517 [0.464, 0.572] | 0.474 [0.422, 0.527] |
| full variable domain | Full heavy chain | 0.508 [0.460, 0.559] | **0.522 [0.472, 0.570]** | 0.514 [0.470, 0.551] |

† VJ-clonotype and V-clonotype are two-stage clonotype retrieval (§2.3), not input representations: reference candidates are restricted to the query's germline gene (V *and* J for VJ-clonotype; V only for V-clonotype), then ranked by CDR3 similarity, and a query with no same-gene reference is scored as a miss. The exact-clonotype filter (VJ-clonotype) is strict and falls below CDR3; relaxing to V only (V-clonotype) recovers and exceeds CDR3. Once the gene is fixed the methods converge (here R@1 ≈ 0.40 at VJ-clonotype for all three). The same ordering holds on VDJdb TCR, McPAS and SAbDab (SAbDab V/J assigned by ANARCI; Supplementary Table S1).

The paratope → framework-CDR3 control → full variable domain progression isolates: (i) the paratope → framework-CDR3 control step, which swaps the native CDR1/CDR2 loops for glycine while adding the framework (+2.9 pp ESM2-150M but -2.1 pp BLOSUM62 and -4.1 pp Levenshtein); and (ii) the clean, length-matched contribution of native CDR1/CDR2 amino-acid content (framework-CDR3 control → full variable domain, +4.0 pp ESM2-150M, +0.5 pp BLOSUM62). For the PLM the two steps are comparable in size, so neither the framework alone nor native CDR1/CDR2 identity alone dominates the paratope→full variable domain gain (and because the paratope→framework-CDR3 control step changes two things at once, we read it descriptively); the alignment methods already reach their ladder peak at paratope (all CDRs) and gain nothing from framework context.

***Table S4. TOST equivalence test for BCR full variable domain (ESM2-150M vs. BLOSUM62).*** *Two one-sided tests on the per-seed Recall@1 differences* $d_i$ = *R@1(ESM2-150M)*$_i$ *- R@1(BLOSUM62)*$_i$ *across the 20 canonical seeds (per-seed values are available in the released data files). Across all ten method pairs at full variable domain, nine are equivalent at Δ = 0.05; the sole exception is ESM2-650M vs. Levenshtein (per-seed mean difference +0.034, 90% CI crossing the margin), i.e. the best-performing ESM2 model (ESM2-650M) sits marginally above the weakest alignment method.*

| Statistic | Value |
|---|---|
| n (seeds) | 20 |
| d_mean | -0.0054 R@1 |
| d_SD | 0.0764 |
| d_SE | 0.0171 |
| Equivalence margin Δ | 0.05 |
| $t_1$ / $t_2$ | +2.61 / -3.24 |
| t_crit (α = 0.05) | ± 1.729 |
| $p_1$ / $p_2$ | ≈ 0.009 / ≈ 0.002 |
| Equivalent at α = 0.05 (Δ = 0.05)? | **Yes** |

***Table S5. Paired-chain retrieval (20 seeds). (A) TCR alpha–beta (VDJdb full variable domain).*** *Single-chain queries use the beta chain only; paired-chain queries use full-length alpha + beta chains, on an identical clone-aware split. TCRdist uses each chain's V-gene-derived CDR1/CDR2/CDR2.5 plus CDR3 via the* `tcrdist3` *package. TCRdist significantly exceeds BLOSUM62 and ESM2-150M on paired chains (non-overlapping 95% CIs); BLOSUM62, Levenshtein and ESM2-150M are mutually comparable.* ***(B) Antibody heavy–light pairing.*** *Paired heavy–light Recall@1 [95% CI] on a small exploratory IEDB set (~45 human, 12 antigens) and on SAbDab heavy–light pairs. Within each dataset all five benchmark methods are statistically indistinguishable (broadly overlapping CIs); SAbDab sits at a lower absolute level, consistent with its lower baseline at every representation. Numbers are nested-bootstrap means.*

*(A) TCR alpha–beta (VDJdb full variable domain)*

| Method | Single-chain mean R@1 | Paired-chain mean R@1 | Δ (paired - single) |
|---|---|---|---|
| Levenshtein | 0.378 | 0.436 | +0.058 |
| BLOSUM62 | 0.399 | 0.435 | +0.037 |
| TCRdist (`tcrdist3`) | 0.428 | **0.515** | +0.087 |
| ESM2-150M | 0.390 | 0.423 | +0.033 |

***Table S5. Paired-chain retrieval (20 seeds). (A) TCR alpha–beta (VDJdb full variable domain).*** *Single-chain queries use the beta chain only; paired-chain queries use full-length alpha + beta chains, on an identical clone-aware split. TCRdist uses each chain's V-gene-derived CDR1/CDR2/CDR2.5 plus CDR3 via the `tcrdist3` package. TCRdist significantly exceeds BLOSUM62 and ESM2-150M on paired chains (non-overlapping 95% CIs); BLOSUM62, Levenshtein and ESM2-150M are mutually comparable.* ***(B) Antibody heavy–light pairing.*** *Paired heavy–light Recall@1 [95% CI] on a small exploratory IEDB set (~45 human, 12 antigens) and on SAbDab heavy–light pairs. Within each dataset all five benchmark methods are statistically indistinguishable (broadly overlapping CIs); SAbDab sits at a lower absolute level, consistent with its lower baseline at every representation. Numbers are nested-bootstrap means.*

*(B) Antibody heavy–light paired R@1 [95% CI]*

| Method | IEDB (~45) | SAbDab |
|---|---|---|
| Levenshtein | 0.533 [0.459, 0.605] | 0.357 [0.289, 0.442] |
| BLOSUM62 | 0.532 [0.457, 0.601] | 0.360 [0.297, 0.450] |
| ESM2-150M | 0.539 [0.468, 0.608] | 0.352 [0.281, 0.442] |
| ESM2-650M | 0.523 [0.453, 0.596] | 0.353 [0.282, 0.433] |
| ESM2-3B | 0.519 [0.445, 0.589] | 0.355 [0.290, 0.438] |

***Table S6. Clone-aware threshold sensitivity (BCR, 20-seed means).*** *The CDR3-similarity clone threshold was varied across six values; expected Recall@1 (mean across the 20 canonical seeds) is reported for BLOSUM62 and ESM2-150M at CDR3 and full variable domain (`threshold_sweep_tie_v2.csv`; matches Supplementary Figure S4).* ***Bold*** *row marks the canonical threshold (0.95). Absolute R@1 is lower at stricter (lower) thresholds, as more near-duplicate leakage is removed; the CDR3 advantage of BLOSUM62 over ESM2-150M (10–13 pp) holds at every threshold, and the two are comparable at full variable domain. Per-seed means here differ from the nested-bootstrap means in Table 1 by <1 pp.*

| Threshold | BLOSUM62 CDR3 | ESM2-150M CDR3 | Δ CDR3 | BLOSUM62 full variable domain | ESM2-150M full variable domain |
|---|---|---|---|---|---|
| 0.85 | 0.451 | 0.325 | +0.126 | 0.394 | 0.396 |
| 0.90 | 0.456 | 0.323 | +0.133 | 0.438 | 0.410 |
| 0.93 | 0.455 | 0.348 | +0.107 | 0.515 | 0.500 |

| Threshold | BLOSUM62 CDR3 | ESM2-150M CDR3 | Δ CDR3 | BLOSUM62 full variable domain | ESM2-150M full variable domain |
|---|---|---|---|---|---|
| **0.95** | **0.455** | **0.353** | +0.102 | **0.513** | **0.507** |
| 0.97 | 0.488 | 0.389 | +0.100 | 0.525 | 0.512 |
| 0.99 | 0.488 | 0.389 | +0.100 | 0.525 | 0.512 |

***Table S7. Per-antigen Recall@1 at BCR CDR3 (CDR3-only), mean across 20 seeds****, for the five adequately-powered antigens. Five rare antigens (N_test/seed < 5 at full variable domain: Ebola GP, RSV F, Spike[IBV], Spike[Rodent CoV], Spike[Bat HKU4]) are omitted as underpowered; the alignment-over-PLM ordering holds among them as well. The micro-averaged BCR CDR3 (R@1 pooled over all test queries, i.e. query-weighted) is reported in main-text Table 1.*

| Antigen | Levenshtein | BLOSUM62 | ESM2-150M | ESM2-650M | ESM2-3B |
|---|---|---|---|---|---|
| HCV polyprotein | 0.480 | 0.442 | 0.419 | 0.420 | 0.440 |
| HIV-1 envelope | 0.477 | 0.485 | 0.330 | 0.381 | 0.322 |
| Influenza HA | 0.460 | 0.485 | 0.409 | 0.381 | 0.382 |
| *Plasmodium* CSP | 0.491 | 0.488 | 0.371 | 0.388 | 0.425 |
| SARS-CoV-2 spike | 0.417 | 0.461 | 0.323 | 0.345 | 0.304 |

***Table S8. Per-antigen Recall@1 at BCR full variable domain (full heavy chain), mean ± SD across 20 seeds****, for the five antigens with N_test/seed ≥ 5. Five underpowered antigens (N_test/seed < 5: Ebola GP, Spike[IBV], RSV F, Spike[Rodent CoV], Spike[Bat HKU4]) are omitted from the rows; the Global micro-average is computed over all antigens.*

| Antigen | N_test/seed | BLOSUM62 R@1 | ESM2-150M R@1 | Δ (ESM2 - BLOSUM62) |
|---|---|---|---|---|
| SARS-CoV-2 spike | 30.2 | 0.542 ± 0.131 | 0.447 ± 0.113 | -0.095 |
| *Plasmodium* CSP | 29.1 | 0.804 ± 0.164 | 0.703 ± 0.181 | -0.100 |
| HIV-1 envelope | 27.8 | 0.449 ± 0.209 | 0.661 ± 0.112 | +0.212 |
| Influenza HA | 20.4 | 0.433 ± 0.123 | 0.361 ± 0.121 | -0.072 |
| HCV polyprotein | 8.2 | 0.309 ± 0.171 | 0.295 ± 0.215 | -0.013 |
| **Global micro-average** | **128.9** | **0.513** | **0.507** | -0.005 |

***Table S9. SHM-stratified BCR full variable domain Recall@1.*** *SHM density is defined as the fraction of mismatches in IMGT V-region positions 1–104 relative to the assigned germline allele. Values are mean ± SD across 20 canonical seeds.*

| SHM bin | N_seqs/seed | BLOSUM62 R@1 | ESM2-150M R@1 | Δ (ESM2 - BLOSUM62) |
|---|---|---|---|---|
| Low (<5%) | 19.5 | 0.586 ± 0.163 | 0.496 ± 0.105 | -0.090 |
| Medium (5– | 70.3 | 0.503 ± 0.163 | 0.485 ± 0.137 | -0.018 |

| SHM bin | N_seqs/seed | BLOSUM62 R@1 | ESM2-150M R@1 | Δ (ESM2 - BLOSUM62) |
|---|---|---|---|---|
| 15%) | | | | |
| High (≥ 15%) | 39.1 | 0.459 ± 0.119 | 0.529 ± 0.104 | +0.070 |

The three bins sum to ~128.6 sequences/seed; the remaining ~3/seed lack an assignable germline allele and are unbinned. Overall BCR full variable domain R@1 (per-seed means, BLOSUM62 / ESM2-150M = 0.513 / 0.507, matching the canonical-threshold row of Table S6; the nested-bootstrap micro-average is in Table S8). SHM density and antigen identity are correlated in IEDB data: the High SHM bin is substantially composed of HIV-1 envelope sequences (~ 63%), so the crossover should be read with the antigen-confounding caveat (Note 2.3).

***Table S10. BLAST CDR3 Recall@1 across four datasets, with BLOSUM62 and ESM2-150M cross-reference.*** *BLAST is included as a second independent alignment baseline; the directional finding that alignment outperforms PLMs is reproduced on all four datasets. Nested-bootstrap 95% CI in brackets.*

| Dataset CDR3 | BLAST R@1 | BLOSUM62 R@1 | ESM2-150M R@1 |
|---|---|---|---|
| BCR (IEDB) | 0.430 [0.387, 0.477] | 0.459 [0.421, 0.499] | 0.357 [0.320, 0.392] |
| VDJdb TCR | 0.361 [0.337, 0.385] | 0.376 [0.352, 0.399] | 0.282 [0.260, 0.305] |
| SAbDab | 0.340 [0.274, 0.423] | 0.362 [0.286, 0.452] | 0.288 [0.228, 0.360] |
| McPAS-TCR | 0.381 [0.360, 0.403] | 0.387 [0.364, 0.408] | 0.272 [0.254, 0.290] |

BLAST tracks just below BLOSUM62 on all four datasets (by 0.6–2.9 pp, including only 0.6 pp on McPAS-TCR) and above every ESM2-150M result; both alignment-based baselines behave similarly at CDR3-only representation, and BLAST's independent seed-extension local alignment corroborates the alignment-over-PLM advantage. Recall@1 is the order-independent expected value under uniform tie resolution, identical in definition to the BLOSUM62/ESM2-150M columns. BLAST seed counts and dataset sizes match the canonical Grand Slam V2 protocol (20 seeds for all four databases; VDJdb TCR uses 10 working epitope labels; n_train/n_test per seed identical to BLOSUM62/ESM2-150M runs; BCR n_train ≈ 546, n_test ≈ 128).

***Table S11. V-gene family sharing in top-1 retrievals (BCR).*** *Fraction of top-1 retrieved query–reference pairs sharing the same V-gene family. The CDR3 row contrasts the substantially lower same-V rate at CDR3-only representation with the full variable domain dominance.*

| Representation | Method | Same-V rate (%) | Sequence method R@1 |
|---|---|---|---|
| CDR3 | BLOSUM62 | 63.2 | 0.459 |
| CDR3 | Levenshtein | 60.5 | 0.446 |

| Representation | Method | Same-V rate (%) | Sequence method R@1 |
|---|---|---|---|
| CDR3 | ESM2-150M | 53.7 | 0.357 |
| full variable domain | BLOSUM62 | 99.1 | 0.513 |
| full variable domain | Levenshtein | 99.2 | 0.498 |
| full variable domain | ESM2-150M | 91.5 | 0.507 |
| full variable domain | Two-stage oracle* | 100.0 | 0.507 [0.464, 0.550] |

Two-stage oracle: filter candidates to same V-gene family (Stage 1), then rank by full-length BLOSUM62 (Stage 2). The V-gene-only random baseline (uniform selection among same-V-family candidates) gives full variable domain R@1 = 0.190 [0.181, 0.200]; the random same-V rate is ~ 34.0%. R@1 values in this table are computed on the subset of full variable domain queries with an ANARCI-assignable V-gene allele (the same subset used for the same-family and oracle analyses) and therefore differ by ≤1.1 pp from the full-set full variable domain R@1 in Supplementary Table S1 and main-text Table 1.

***Table S12. Benchmark representations: what each isolates and what its result indicates.*** *The ladder is designed so that each step changes as little as possible (the framework-CDR3 control substitution control is the exception: it both adds the framework and masks CDR1/CDR2; see that row) and thereby tests one claim at a time. CDR3, paratope and full variable domain vary the input along a germline axis (junctional CDR3 → all antigen-contact loops → full chain including germline framework); VJ- and V-clonotype hold the CDR3 input fixed but change the retrieval procedure (add a germline-gene filter); framework-CDR3 control is a length-matched control that adds the framework while masking native CDR1/CDR2. Numbers are 20-seed nested-bootstrap Recall@1 (BCR unless noted); see Table S3, Supplementary Table S1, and §3.2–3.3.*

| Representation | Input / procedure | Key result | What it indicates |
|---|---|---|---|
| **CDR3** | Embed the CDR3 region; rank the whole reference DB | Alignment beats every ESM2 size by 7–12 pp on all four DBs (BLOSUM62 0.459 vs. ESM2-150M 0.357) | The antigen-discriminative signal is concentrated in the hypervariable, non-germline CDR3, and simple alignment captures it better than zero-shot PLM embeddings. |
| **VJ-clonotype** (V+J filter) | Restrict candidates to the same V *and* J gene, then rank by CDR3; miss if none | All seven methods converge to R@1 ≈ 0.40 (BCR 0.39–0.43); falls *below* CDR3; paired chains | Once retrieval is conditioned on germline-gene identity, the |

| Representation | Input / procedure | Key result | What it indicates |
|---|---|---|---|
| | | converge to within 0.001 (0.122–0.123) | choice of CDR3 comparison becomes irrelevant: the germline-gene filter, not the method, carries the retrievable signal. |
| **V-clonotype** (V filter) | As VJ-clonotype but require only a shared V gene | Recovers above CDR3 (BCR BLOSUM62 0.526, ESM2-150M 0.483); alignment regains a small lead | VJ-clonotype's strictness is driven by the J-gene requirement; relaxing it enlarges the same-gene pool. The residual small alignment lead is a milder form of the CDR3 effect. |
| **paratope** (All CDRs) | Embed CDR1+CDR2+CDR3 concatenated | BLOSUM62 0.538, ESM2-150M 0.445; gap ~9 pp, similar to CDR3 | Adding CDR1 and CDR2 helps both families by about the same amount (ESM2-150M +8.8 pp, BLOSUM62 +7.9 pp over CDR3): the germline-derived loops carry a method-neutral signal, so the CDR3 ordering simply carries through. |
| **framework-CDR3 control** (Framework-only control) | CDR1/CDR2 replaced by length-matched Gly spacers; FW1–FW4 + native CDR3 retained | ESM2-150M: paratope→framework-CDR3 control (adds framework, glycine-masks CDR1/2) +2.9 pp; framework-CDR3 control→full variable domain (restores native | The PLM's full-length gain draws on both the germline framework and the germline-derived |

| Representation | Input / procedure | Key result | What it indicates |
|---|---|---|---|
| | | CDR1/2) +4.0 pp, comparable steps, read descriptively since paratope→framework-CDR3 control changes two things at once; alignment gains little from the framework | CDR1/CDR2 loops, consistent with a germline shortcut rather than learned CDR-level antigen specificity. |
| **Full variable domain** | Embed the full variable domain (framework + all CDRs) | The five benchmark methods cluster within ~3.3 pp (Levenshtein 0.508 to ESM2-650M 0.541; best PLM now marginally above best alignment) | Adding the germline framework brings PLM and alignment to parity: the convergence is best read as a framework-associated *germline shortcut* rather than learned CDR-level antigen specificity. |

***Table S13. Secondary retrieval metrics (Recall@5 and MRR) reproduce the Recall@1 ranking.*** *Twenty-seed nested-bootstrap means of expected Recall@5 and mean reciprocal rank (MRR) for the five benchmark methods, at the CDR3 input (CDR3) and full chain (full variable domain), across all four databases. At CDR3, BLOSUM62 leads on both secondary metrics on every database (exactly as for Recall@1), and at full variable domain the leadership scatters across methods within a narrow band, again mirroring the Recall@1 convergence. The choice of Recall@1 as the primary metric is therefore not what drives the method ranking. Bold marks the highest value in each row. Values are the 20-seed post-filter nested-bootstrap means (same canonical source as Table S1).*

| Database | Level | Metric | Levenshtein | BLOSUM62 | ESM2-150M | ESM2-650M | ESM2-3B |
|---|---|---|---|---|---|---|---|
| BCR (IEDB) | CDR3 | R@5 | 0.701 | **0.717** | 0.651 | 0.700 | 0.661 |
| | CDR3 | MRR | 0.562 | **0.575** | 0.488 | 0.510 | 0.487 |
| | full variable domain† | R@5 | 0.765 | 0.763 | **0.801** | 0.794 | 0.793 |
| | full variable domain† | MRR | 0.619 | 0.629 | 0.634 | **0.653** | 0.641 |

| Database | Level | Metric | Levenshtein | BLOSUM62 | ESM2-150M | ESM2-650M | ESM2-3B |
|---|---|---|---|---|---|---|---|
| VDJdb TCR | CDR3 | R@5 | 0.613 | **0.619** | 0.563 | 0.557 | 0.574 |
| | CDR3 | MRR | 0.478 | **0.491** | 0.420 | 0.431 | 0.437 |
| | full variable domain‡ | R@5 | 0.630 | 0.629 | 0.611 | 0.620 | **0.633** |
| | full variable domain‡ | MRR | 0.501 | **0.511** | 0.501 | 0.505 | 0.507 |
| SAbDab | CDR3 | R@5 | 0.558 | **0.564** | 0.508 | 0.509 | 0.506 |
| | CDR3 | MRR | 0.440 | **0.446** | 0.380 | 0.374 | 0.368 |
| | full variable domain† | R@5 | 0.515 | 0.534 | 0.537 | 0.545 | **0.547** |
| | full variable domain† | MRR | 0.425 | 0.444 | **0.448** | 0.411 | 0.444 |
| McPAS-TCR | CDR3 | R@5 | 0.618 | **0.643** | 0.582 | 0.584 | 0.595 |
| | CDR3 | MRR | 0.484 | **0.508** | 0.418 | 0.427 | 0.442 |
| | full variable domain‡ | R@5 | 0.703 | 0.708 | **0.713** | 0.702 | 0.702 |
| | full variable domain‡ | MRR | 0.553 | 0.555 | **0.557** | 0.544 | 0.544 |

† Real full-length chains (BCR heavy chain; SAbDab paired heavy+light). ‡ VDJdb and McPAS-TCR full variable domain use germline-reconstructed full chains and are directional (main text §2.2), as in Table 1.

***Table S14. SCEPTR vs. alignment vs. `tcrdist3` on the common gene-mappable query set (main text §3.5).*** *Expected Recall@1 (20-seed means) for all five methods scored on the identical per-seed clone-aware split and restricted to the common subset of test queries whose V/J genes `tcrdist3` can map (kept fraction shown). SCEPTR embeddings were generated in an isolated environment and scored through the same pipeline as every other method (Supplementary Methods, SCEPTR; Common gene-mappable comparison set). Because this gene-mappable subset differs slightly from the full test set, values differ modestly from the canonical full-set numbers in Table S1 (e.g. VDJdb single-chain BLOSUM62 0.311 here versus 0.375 full-set; this is also why the common-subset `tcrdist3` paired value 0.490 differs from the full-set 0.515 of Table S5). Bold marks the highest value in each row. Note on the CDR3 condition: unlike BLOSUM62, Levenshtein and ESM2-150M, which see only the CDR3 amino-acid string, both `tcrdist3` and SCEPTR incorporate the V gene even here (`tcrdist3` scores the V-gene-derived CDR1/CDR2/CDR2.5 loops alongside*

*CDR3; SCEPTR encodes the V gene in its embedding). Their CDR3-level edge is therefore the germline V-gene signal that CDR3-only methods lack, and it vanishes once alignment is given the same germline context through the clonotype filter (main text §3.5). SCEPTR is thus the only language model to reach alignment-level CDR3 retrieval, matching the germline-informed `tcrdist3` at single-chain CDR3 but not on paired chains, where only `tcrdist3` leads.*

| Dataset (input) | Kept | `tcrdist3` | SCEPTR | BLOSUM62 | Levenshtein | ESM2-150M |
|---|---|---|---|---|---|---|
| McPAS-TCR (CDR3) | 198/199 | **0.463** | 0.457 | 0.393 | 0.365 | 0.275 |
| VDJdb (CDR3) | 253/272 | **0.338** | 0.333 | 0.311 | 0.280 | 0.219 |
| VDJdb (paired α+β) | 113/199 | **0.490** | 0.468 | 0.467 | 0.449 | 0.389 |

***Table S15. Domain-specific and auxiliary models (companion to Table S1).*** *Expected Recall@1 (20-seed nested-bootstrap means) for the antibody-specific (AbLang, AntiBERTy), TCR-specific (TCR-BERT, SCEPTR) and newer general-purpose (ESM-C-300m) models, by database and receptor representation. "—" marks a representation not assessed for that model: AbLang/AntiBERTy are antibody-specific (BCR, SAbDab) and TCR-BERT/SCEPTR are TCR-specific (VDJdb, McPAS); SCEPTR is a fixed V+CDR3 model with no paratope or full-domain input, and TCR-BERT's 64-token limit excludes the full domain (ESM-C-300m, embedded externally on GPU, is reported at all representations). *The TCR-BERT paratope is out-of-distribution (the model is trained on CDR3 alone) but is reported for completeness: it still gains from the germline-encoded CDR1/CDR2 loops, landing above ESM2-150M but below alignment, consistent with the germline shortcut. SCEPTR's single-chain CDR3 head-to-head against `tcrdist3` on the common gene-mappable subset is in Table S14; the purpose-built `tcrdist3` and the borrowed BCRdist control are summarized in the footnote of Table S1. Full 95% CIs for every cell are in the released `Table_S1_Full_CI.csv`.*

| Model | Database | CDR3 | VJ-clono | V-clono | Paratope | Full domain |
|---|---|---|---|---|---|---|
| AbLang | BCR | 0.390 | 0.400 | 0.479 | 0.504 | 0.535 |
| AbLang | SAbDab | 0.322 | 0.258 | 0.323 | 0.336 | 0.341 |
| AntiBERTy | BCR | 0.398 | 0.428 | 0.495 | 0.471 | 0.551 |
| AntiBERTy | SAbDab | 0.286 | 0.251 | 0.314 | 0.325 | 0.373 |
| TCR-BERT | VDJdb | 0.349 | 0.305 | 0.334 | 0.387* | — |
| TCR-BERT | McPAS | 0.333 | 0.384 | 0.471 | 0.424* | — |
| SCEPTR | VDJdb | 0.404 | 0.306 | 0.350 | — | — |
| SCEPTR | McPAS | 0.450 | 0.393 | 0.482 | — | — |
| ESM-C-300m | BCR | 0.387 | 0.419 | 0.488 | 0.376 | 0.499 |
| ESM-C-300m | VDJdb | 0.284 | 0.295 | 0.327 | 0.359 | 0.386 |

| Model | Database | CDR3 | VJ-clono | V-clono | Paratope | Full domain |
|---|---|---|---|---|---|---|
| ESM-C-300m | SAbDab | 0.287 | 0.252 | 0.294 | 0.245 | 0.321 |
| ESM-C-300m | McPAS | 0.288 | 0.378 | 0.461 | 0.388 | 0.435 |

### 4 Supplementary Figures

This section reports eleven supplementary figures (S1–S11) supporting the main paper Results (§3) and Supplementary Notes (§S2.1–S2.6).

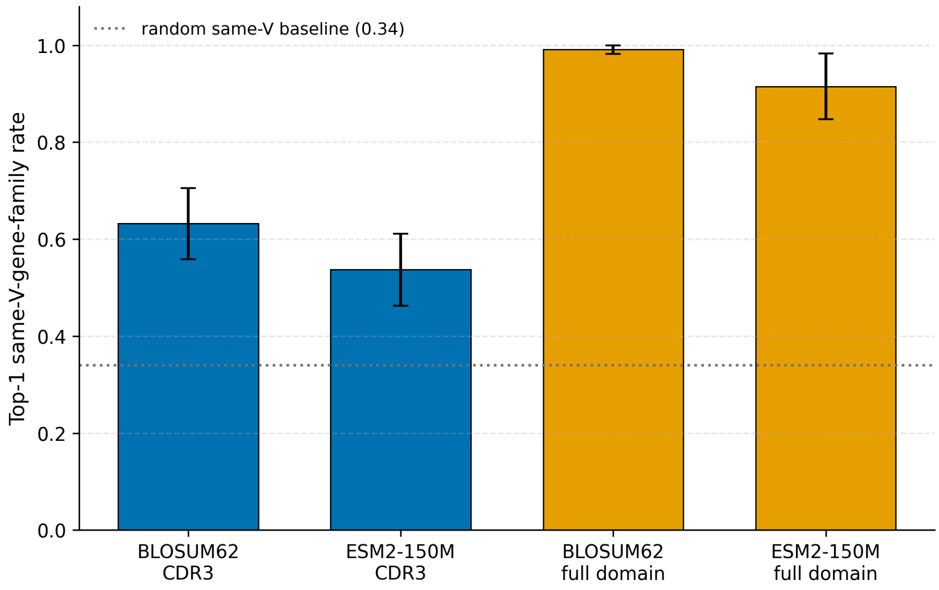


***Figure S1.*** *V-gene family agreement between query and top-1 retrieved candidate at BCR CDR3 (blue) and full variable domain (orange), for BLOSUM62 and ESM2-150M. The fraction of retrievals sharing the query's V-gene family rises from ~0.6 at CDR3 (BLOSUM62 0.63, ESM2-150M 0.54) to ~0.95 at full variable domain (BLOSUM62 0.99, ESM2-150M 0.92), far above the random same-family baseline (dotted line, 0.34; observed-to-random ratios 1.83×/1.56× at CDR3, 2.92×/2.70× at full variable domain). This substantial CDR3→full variable domain increase is the signature we summarize as the germline shortcut. Error bars: SD across 20 seeds.*

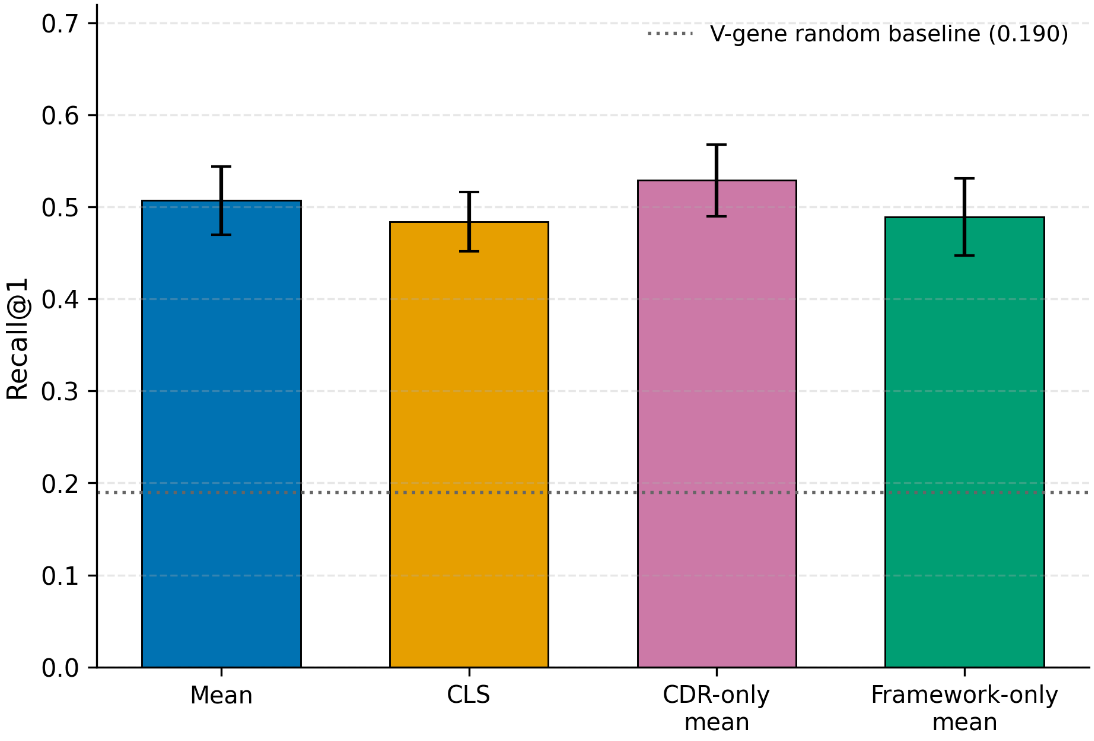


***Figure S2.*** *Pooling sensitivity on cleaned variable-domain inputs. ESM2-150M BCR full variable domain Recall@1 for four pooling strategies: mean (0.507), CLS (0.484), CDR-only mean (average over CDR1/CDR2/CDR3 tokens; 0.529) and framework-only mean (average over variable-domain non-CDR tokens; 0.489). All four are statistically indistinguishable (overlapping 95% CIs) and CLS shows no collapse; the dotted horizontal line is the V-gene-only random baseline (R@1 = 0.190), far below all four. The full variable domain result is therefore robust to pooling choice. Because global self-attention means every retained token has already attended to the rest of the chain, neither region-restricted pooling isolates its region's signal; the only directional difference (CDR-only marginally above framework-only) is the opposite of what a framework-localized signal would predict, so we read this figure as a robustness / no-collapse check rather than as localization evidence (Note S2.2). Error bars: 95% CI.*

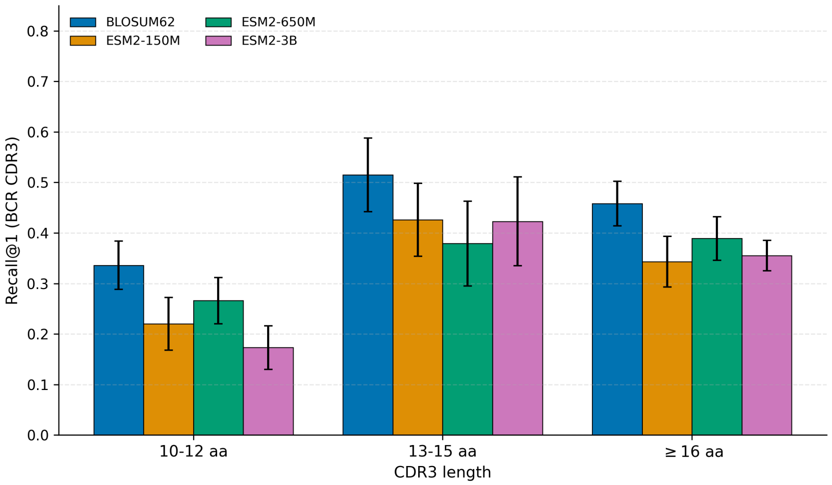


***Figure S3. CDR3 length stratification.*** *Per-length-bin BCR CDR3 Recall@1 for BLOSUM62, ESM2-150M, ESM2-650M and ESM2-3B across three CDR3 length buckets (10–12, 13–15, ≥ 16 aa). The alignment advantage persists across the CDR3 length distribution and is not driven by short or long CDR3 sub-populations. Error bars: 95% CI.*

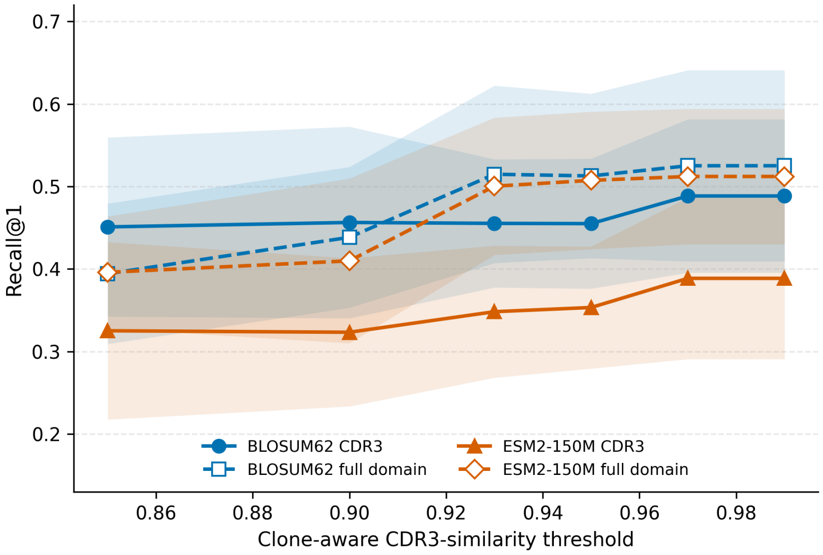


***Figure S4. Clone-threshold robustness (cleaned BCR set).*** *BCR CDR3 and full variable domain Recall@1 for BLOSUM62 and ESM2-150M across CDR3-similarity clone-split thresholds (0.85–0.99; 20 seeds). Retrieval changes slightly across thresholds: approximately 12% of the CDR3 sequences have neighbors in the [0.85, 0.99) similarity range, causing minor connected-component split fluctuations and leading to small variations in expected R@1 (maximum CDR3 variation across thresholds ~6.6 pp for ESM2-150M and ~3.7 pp for BLOSUM62). Nevertheless, the qualitative conclusions are fully robust: BLOSUM62 leads ESM2-150M at CDR3 (CDR3) across all thresholds, while at full variable domain (full chain) the two are comparable, consistent with the main-text results. Error bars: SD across 20 seeds.*

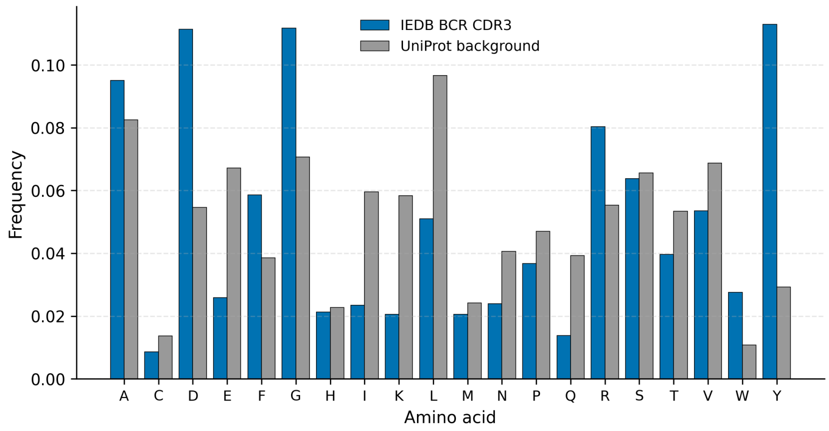


***Figure S5. IEDB BCR CDR3 amino-acid composition vs. UniProt background.*** *Per-residue frequency in the IEDB BCR CDR3 benchmark sequences compared with the UniProt protein-wide background frequency. BCR CDR3 are enriched in Y, D, A and G and depleted in L, E, I; this compositional difference is biological (V(D)J junctional residues) and does not introduce systematic bias to the retrieval evaluation.*

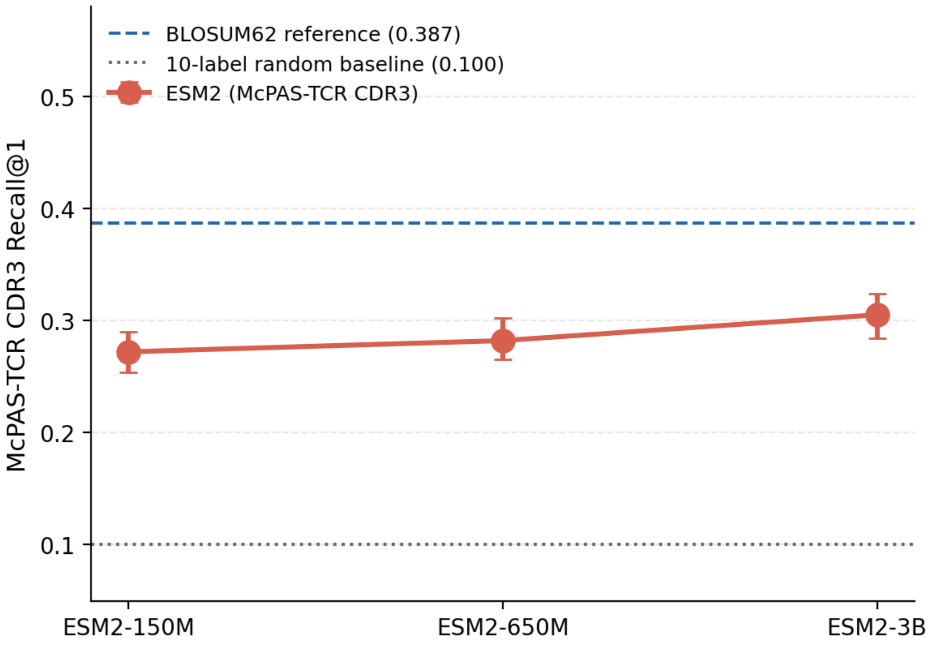


***Figure S6. McPAS-TCR CDR3 (CDR3) retrieval rises modestly with ESM2 scale*** *(expected R@1, 20-seed nested-bootstrap protocol; §2.1). ESM2 Recall@1 increases monotonically across model size, from 0.272 (150M) to 0.282 (650M) to 0.305 (3B), staying above the 10-label random baseline (0.100, dotted) yet well below the BLOSUM62 reference (0.387, dashed). McPAS-TCR is the only one of the four datasets on which ESM2 scales monotonically at CDR3; even at 3B the largest model trails alignment by 8.2 pp, showing that model size alone does not close the CDR3-level gap. Error bars: 95% nested-bootstrap CI.*

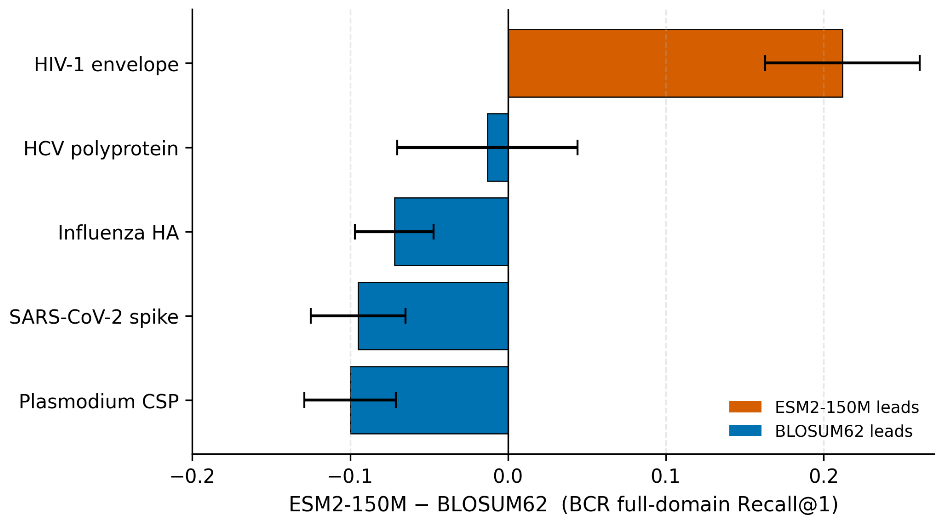


***Figure S7.*** *BCR full variable domain per-antigen method gap: ESM2-150M - BLOSUM62 Recall@1, 20-seed mean ± SE. Antigens are restricted to per-seed N_test ≥ 5. Effect sizes: HIV-1 Env +0.212 ± 0.049, SARS-CoV-2 -0.095 ± 0.030, HCV polyprotein -0.013 ± 0.057, Influenza HA -0.072 ± 0.025, Plasmodium CSP -0.100 ± 0.029. Bars are colored by which method leads (legend); the solid vertical line marks zero (no method gap), and HCV polyprotein straddles it (gap not significant). Five further antigens with per-seed N_test < 5 are omitted from both the figure and Table S8 (included only in the latter's global micro-average). HIV-1 envelope is the only major antigen where the PLM leads, and is the largest single-antigen divergence; it motivates the SHM-stratified analysis (Note 2.3).*

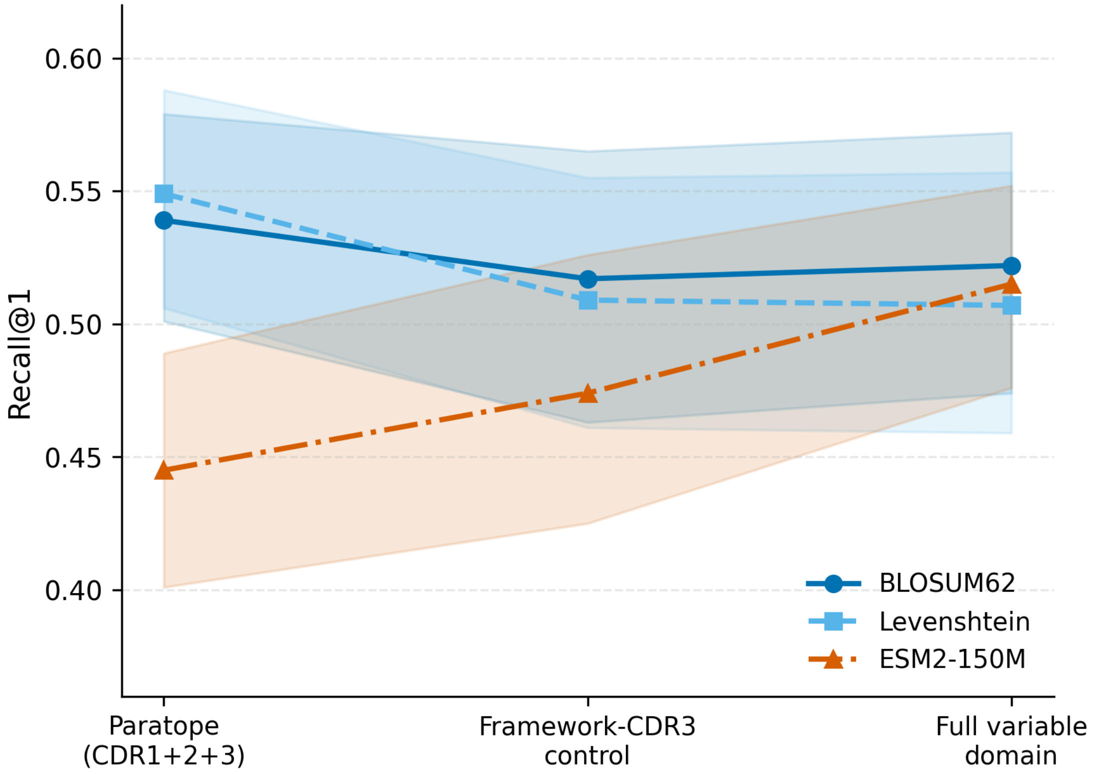


***Figure S8.*** *Framework-CDR3 control isolating the contribution of CDR1/CDR2 amino-acid content. Three-point trajectory from paratope (CDR1+CDR2+CDR3, no framework) through framework-CDR3 control (length-matched framework only: FW1+G_|CDR1|+FW2+G_|CDR2|+FW3+CDR3+FW4) to full variable domain (full variable region). The paratope → framework-CDR3 control step (add framework, mask CDR1/CDR2) is +2.9 pp for ESM2-150M but -2.1 pp for BLOSUM62; the clean length-matched framework-CDR3 control → full variable domain step quantifies the contribution of native CDR1/CDR2 amino-acid identity (+4.0 pp for ESM2-150M, +0.5 pp for BLOSUM62). The two ESM2-150M steps are comparable in size, so the paratope→full variable domain recovery is dominated by neither the framework alone nor native CDR1/CDR2 identity alone; because the paratope→framework-CDR3 control step changes two things at once (adds framework, masks CDR1/2), we read the step sizes descriptively rather than as a clean framework contribution. BLOSUM62 already extracted the available signal from the CDRs alone and gains nothing from the framework. Error bars: 95% nested-bootstrap CI across 20 seeds.*

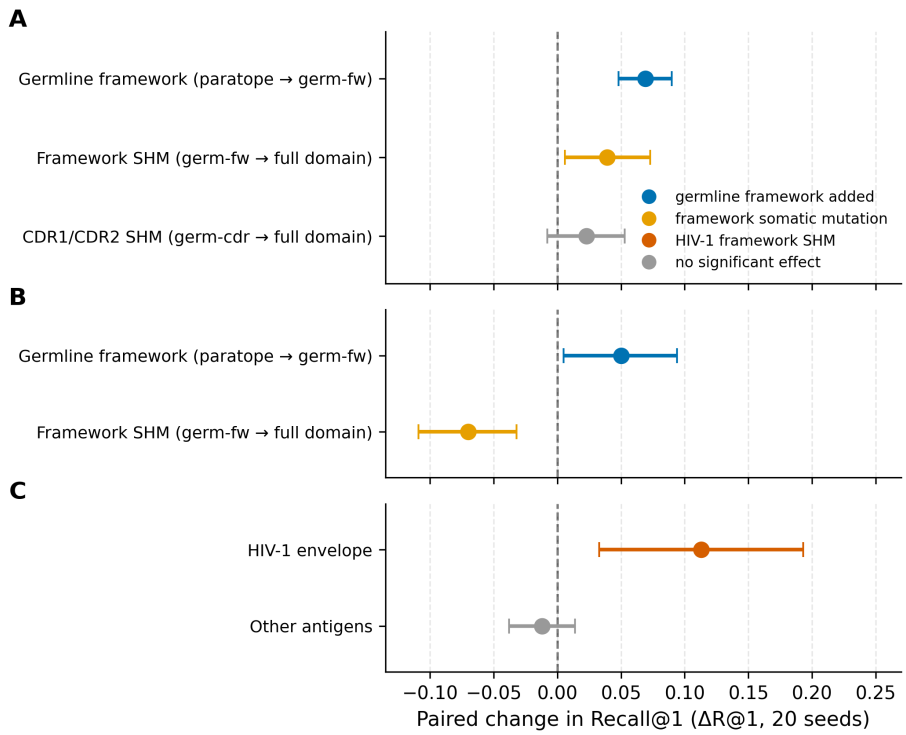


***Figure S9.*** *Germline-reversion test: where the PLM's full-length gain comes from. Per-seed paired change in Recall@1 (ESM2-150M, 20 seeds) when one region of each antibody is reverted to its assigned germline IGHV sequence while the rest stays native (Note S2.5).* ***(A)*** *BCR (ESM2-150M): adding the germline framework to the all-CDR input (paratope → germ-fw) recovers most of the full-length gain, whereas the framework's own somatic mutations (germ-fw → full variable domain) and CDR1/CDR2 somatic mutations (germ-cdr → full variable domain) add little.* ***(B)*** *SAbDab (ESM2-150M): the germline-framework gain replicates on a second native antibody database, but native framework mutations there reduce retrieval (its structure-derived, engineered frameworks diverge from germline non-naturally).* ***(C)*** *Framework somatic mutation split by antigen (BCR, after removing pretraining-leak candidates): the effect is confined to HIV-1 envelope (the most hypermutated antibodies) and absent elsewhere. Points are paired mean ΔR@1; bars are 95% confidence intervals; the dashed line marks no change. For BLOSUM62 every step is within noise of zero.*

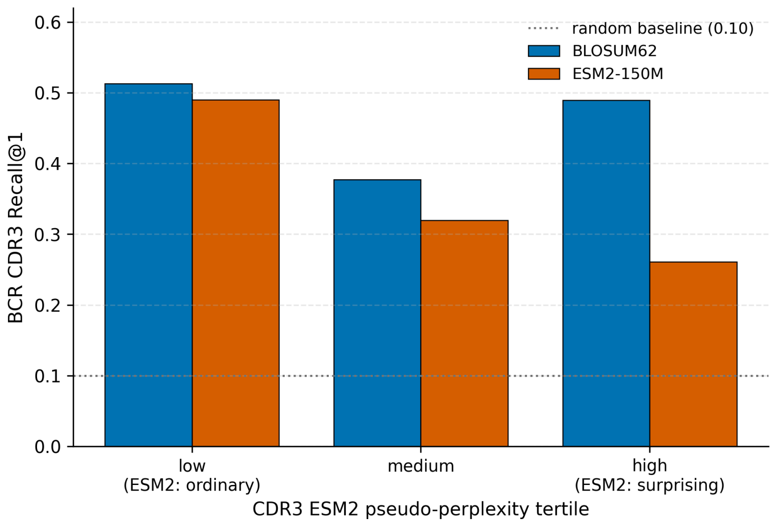


***Figure S10.*** *The PLM's CDR3 deficit is larger on the CDR3s it scores as atypical. Per-query BCR CDR3 Recall@1 for BLOSUM62 and ESM2-150M, binned by the CDR3's ESM2 masked-language pseudo-perplexity (a model-internal per-residue measure of how atypical the CDR3 is to ESM2; low = the model finds it ordinary, high = the model finds it surprising). ESM2-150M retrieval declines monotonically with pseudo-perplexity (0.49, 0.32, 0.26) while BLOSUM62 does not decline monotonically (0.51, 0.38, 0.49; it dips in the medium tertile and recovers in the high tertile): the two are at parity where ESM2 finds the CDR3 ordinary and diverge by 23 pp where it finds it most surprising. Bars pool 20 seeds (2,564 query instances); the deficit-vs-pseudo-perplexity correlation over the 559 unique CDR3s (Spearman -0.14,* $p<10^{-3}$*) is unchanged after controlling for CDR3 length. This pseudo-perplexity is internal to ESM2; a model-independent germline V–D–J distance neither reproduces the gradient nor correlates with pseudo-perplexity (Note S2.6), so the effect is a property of the model's representation, not a measurable junctional/germline divergence. Dotted line: 10-label random baseline.*

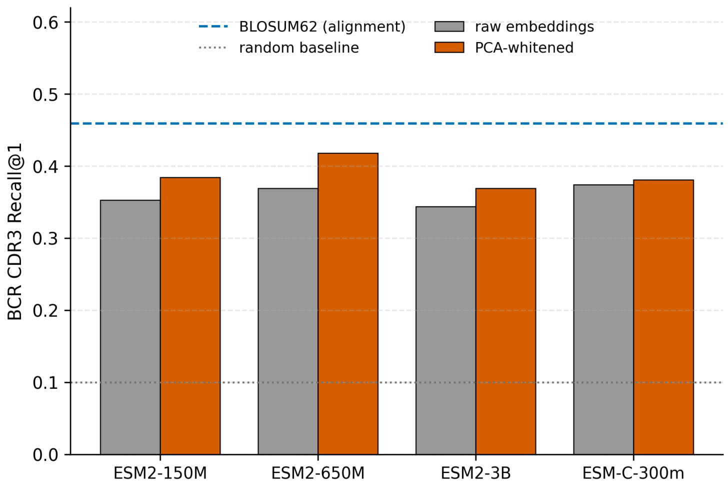


***Figure S11.*** *The CDR3 deficit is representational, not a cosine-geometry artifact (Note S2.6). BCR CDR3 Recall@1 for four zero-shot PLMs, raw versus PCA-whitened embeddings (whitening fit on the reference set only; the single significant isotropy fix). All four PLMs are strongly anisotropic (mean pairwise cosine 0.95–0.99), yet whitening adds only ~3–5 pp and every model stays below the BLOSUM62 alignment baseline (dashed). Two further controls (not shown) rule geometry out directly: the full-chain embeddings, where the PLM is at parity with alignment, are equally anisotropic (0.97–0.99) yet gain nothing from whitening, and the deficit persists under Euclidean distance. 20-seed means (`scratch/cdr3_mechanism_geometry.py`, `cdr3_mechanism_supp.py`); dotted line, 10-label random baseline.*
